# Mapping RNA-Protein Interactions with Subcellular Resolution Using Colocalization CLIP

**DOI:** 10.1101/2023.10.26.563984

**Authors:** Soon Yi, Shashi S. Singh, Kathryn Rozen-Gagnon, Joseph M. Luna

## Abstract

RNA binding proteins (RBPs) are essential for RNA metabolism and profoundly impact health and disease. The subcellular organization of RBP interaction networks with target RNAs remains largely unexplored. Here, we develop colocalization CLIP, a method that combines CrossLinking and ImmunoPrecipitation (CLIP) with proximity labeling, to explore in-depth the subcellular RNA interactions of the well-studied RNA-binding protein HuR. Using this method, we uncover HuR’s dynamic and location-specific interactions with RNA, revealing alterations in sequence preferences and interactions in the nucleus, cytosol, or stress granule compartments. We uncover HuR’s unique binding preferences within stress granules during arsenite stress, illuminating intricate interactions that conventional methodologies cannot capture. Overall, coCLIP provides a powerful method for revealing RBP:RNA interactions based on localization and lays the foundation for an advanced understanding of RBP models that incorporate subcellular location as a critical determinant of their functions.

## Introduction

RNA localization is a critical feature of many cellular processes, including the regulation of gene expression, mRNA stability, and translation. The coordinated efforts of thousands of RNA binding proteins (RBPs) are central in mediating RNA outcomes. These proteins govern nearly all aspects of RNA metabolism, from RNA processing, nuclear export, translation, and eventual turnover.^1^ Despite advancements in sequencing and imaging techniques for RNA localization,^2,3^ pinpointing the exact sites of RBP:RNA interactions in subcellular spaces remains a significant challenge in RNA biology.

Our knowledge of RBP:RNA interactions has been revolutionized in the past decade by CrossLinking and ImmunoPrecipitation (CLIP) methods.^4–8^ These methods employ UV irradiation of cells or tissues to generate covalent bonds between RNA and protein at near-zero distances. These protein:RNA crosslinks enable the stringent purification of a given RBP with its *in vivo* RNA targets for subsequent deep sequencing. Still, CLIP does not preserve spatial information, making it difficult to determine the precise roles of specific RBP:RNA interactions based on localization.

Here, we develop a method to overcome this limitation by combining CLIP with proximity labeling using APEX2^9^ at predefined subcellular locations, termed colocalization CLIP (coCLIP). We establish coCLIP with benchmark tests on HuR, a well-studied RNA binding protein with specific binding profiles and functional roles based on subcellular localization. To set a foundational comparison for coCLIP, we conducted CLIP from nuclear or cytosolic fractions, subsequently employing this technique for side-by-side analysis. HuR coCLIP maintains consistent and specific subcellular binding during arsenite stress compared to CLIP from fractionation. We further focus on the arsenite-induced stress response that results in HuR relocalization to stress granules (SGs). Our results reveal that HuR binding to target RNAs is location-specific and showcases the dynamic nature of HuR-RNA interactions at subcellular resolution. Moreover, HuR coCLIP shows altered sequence preferences for HuR binding within the cytosol and SGs that were previously uncharacterized. Overall, coCLIP provides a powerful method for uncovering RBP:RNA interactions based on localization, revealing how RBP interactions on transcript loci are organized in subcellular spaces.

## Results

### Colocalization CLIP (coCLIP) reveals HuR target interactions at subcellular resolution

To establish a universal method for delineating subcellular RBP:RNA interactions, we integrated CLIP with proximity labeling, utilizing the engineered ascorbate peroxidase APEX2.^9,10^ APEX labeling, and similar proximity-based strategies,^11^ have been previously used to define subcellular proteomes,^12–16^ transcriptomes,^17–19^ and protein-occupied RNAs.^20^ We hypothesized that combining a proximity approach with CLIP would allow us to simultaneously identify global and local RNA targets for specific RBPs (**Figure 1A-B**). We began by engineering cells to express APEX2 at predetermined locations, such as the cytosol, nucleus, or within stress granules, by creating a fusion protein with G3BP1.^21^ After initiating the APEX reaction, we evaluated biotin labeling using fluorescent streptavidin and identified a streptavidin reactive signal primarily confined to the labeling locations, and was stress-dependent in the case of G3BP1-APEX (**Figure 1C, S1A**).

**Figure 1:**
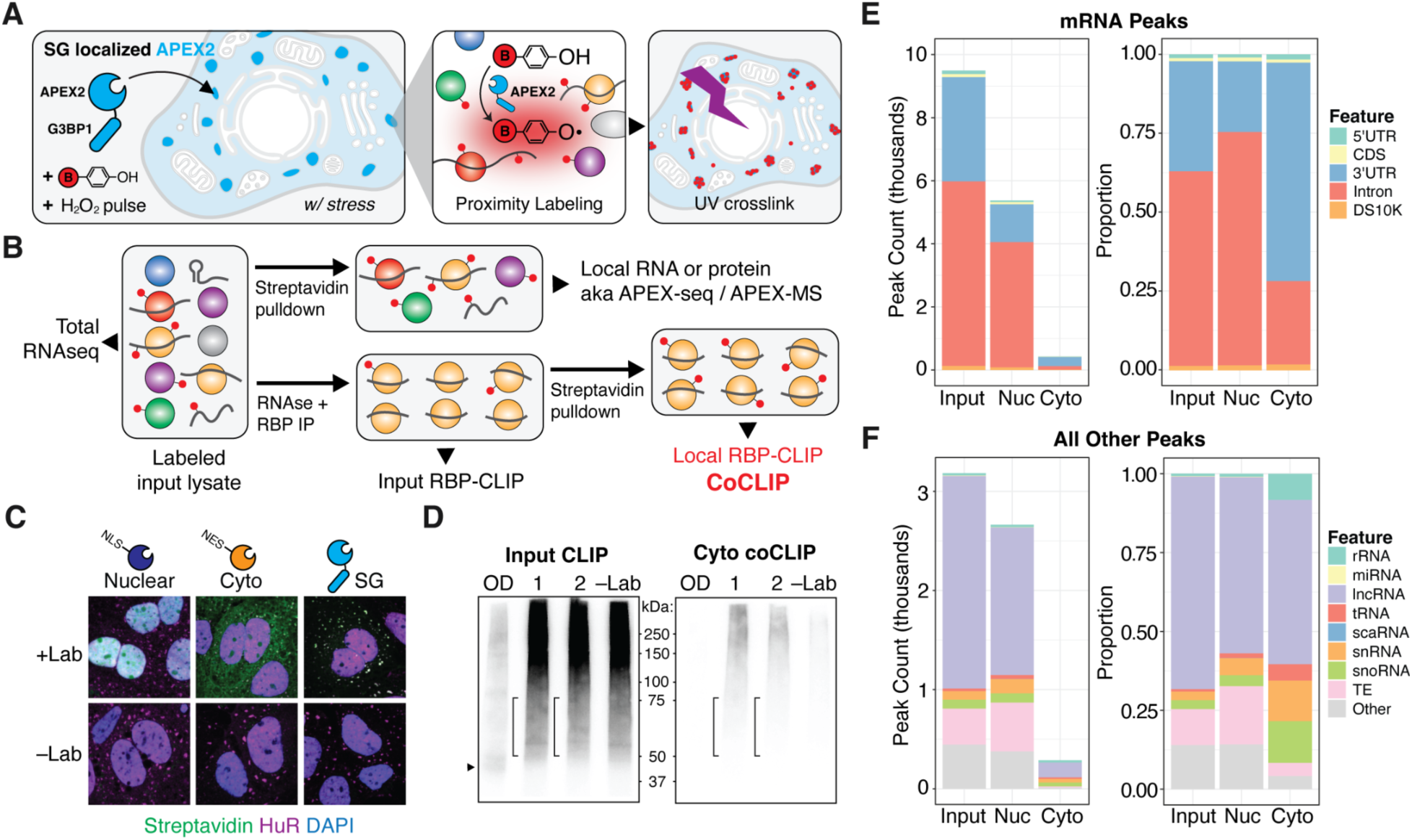
Colocalization CLIP (coCLIP) for subcellular resolution of RBP:RNA interactions. **(A)** An APEX2 enzyme is localized to a specific compartment (depicted here as a G3BP1 fusion for stress granules, SGs). The proximity labeling reaction using biotin phenol (BP) and hydrogen peroxide is performed, followed by UV crosslinking. **(B)** After cell lysis, the sample undergoes streptavidin pulldown to isolate a local proteome or transcriptome or undergoes RBP-CLIP. A second streptavidin pulldown following RBP IP enables the capture of local RBP targets. **(C)** Immunofluorescence for streptavidin reactive signal with and without labeling reagents for arsenite stressed Huh7.5 cells with nuclear, cytoplasmic, or SG localized APEX2. **(D)** Example autoradiograms for input HuR CLIP (left) or cytoplasmic HuR coCLIP (right) from two replicates. Size of HuR indicated with an arrow, excised HuR:RNA complexes in brackets. OD: RNAse overdigestion; -Lab: No label. **(E)** HuR peak counts (left) and proportions (right) on mRNAs for input CLIP, nuclear coCLIP, or cytoplasmic coCLIP in Huh7.5 cells. DS10K: within 10kb downstream of 3’UTRs. **(F)** HuR peak counts (left) and proportions (right) as in **(E)** for ncRNAs. See also Figures S1-S3.

We next focused on HuR as a candidate RBP to assess the viability of our coCLIP approach. HuR is a ubiquitously expressed member of the ELAV family of RNA-binding proteins.^22^ HuR shuttles between the nucleus and cytoplasm and binds to AU-rich motifs in 3’ untranslated regions (3’UTRs) of mRNAs to affect stability.^22^ In the nucleus, HuR also interacts with introns to modulate the alternative splicing of specific mRNAs.^23^ Typically, over 90% of HuR is located in the nucleus; however, under stress conditions, HuR relocates to the cytoplasm and accumulates in SGs,^24^ as illustrated in **Figure S1B**. Given HuR’s ability to interact with different transcript features depending on its location—intronic in the nucleus and 3’UTR in the cytoplasm—it emerged as a suitable candidate RBP for defining subcellular RBP:RNA interactions with coCLIP.

After confirming that UV crosslinking does not induce spurious APEX2 activity or non-specific biotinylation (**Figure S2A**), we initiated APEX labeling in cells, followed immediately by UV crosslinking (see methods). We carried out antibody-based CLIP for HuR in arsenite-stressed and unstressed conditions (**Figure 1D, S2B**) and performed streptavidin enrichment of eluates from HuR pull-downs (**Figure 1D, S2C**). We observed no significant differences in HuR:RNA signal intensities from input HuR-CLIP across samples (**Figure S2B**). However, we noted that HuR coCLIP in the nucleus gave a more robust signal than coCLIP from the cytoplasm or SGs (**Figure S2C**), reflecting the distribution of HuR at steady state.

We performed coCLIP and created sequencing libraries from nuclear, cytosolic, or G3BP tethered APEX2 under mock or sodium arsenite conditions, all with associated input and enriched libraries, for 58 sequenced libraries in total (**Table S1**). To facilitate robust and consistent data analysis, similar to previous efforts for CLIP in non-model organisms,^25^ we further created a custom single command-line analysis pipeline, termed “CLIPpityCLIP,” to process raw sequencing data into peaks for subsequent analysis and visualization (**Figure S3**, see methods). We first compared nuclear and cytoplasmic coCLIP data to input HuR-CLIP from the same samples as a reference, all under unstressed conditions. We retained peaks if they appeared in at least half of the replicates and had reads above the median count across all samples. Input HuR-CLIP yielded over 9000 HuR peaks on mRNAs with a characteristic preference for intronic (60%) and 3’UTR (30%) binding as has been previously reported (**Figure 1E**).^23,26^ Nuclear coCLIP showed enrichment in intronic binding (75%), while cytoplasmic coCLIP revealed a preference for 3’UTR binding (75%) (**Figure 1E**). In general, we observed approximately 10-fold more HuR target peaks from nuclear coCLIP relative to cytoplasmic coCLIP in unstressed cells, reflecting the steady state distribution of HuR within the nucleus versus cytoplasm (**Figure 1E**). We also observed this pattern among non-coding HuR targets, where nuclear coCLIP yielded over 2500 HuR peaks while cytoplasmic coCLIP yielded 10-fold fewer peaks (**Figure 1F**). Taken together, these results suggest that coCLIP can parse dynamic subpopulations of HuR within the nucleus or cytoplasm in situ.

### HuR binding preferences remain consistent with coCLIP during arsenite stress

Next, we aimed to contrast coCLIP with standard CLIP derived from nuclear or cytoplasmic cell fractions. Cellular fractionation is widely used to provide insight into the function and composition of cellular organelles and is particularly useful when studying RBPs,^27,28^ such as HuR,^29^ that undergo nucleocytoplasmic trafficking. Following a widely used protocol,^30,31^ we conducted HuR CLIP from nuclear and cytoplasmic fractions with cells in both arsenite-induced stress and unstressed states (**Figure S4A-B**). As cellular stressors such as arsenite are known to alter nucleocytoplasmic transport,^32^ we reasoned that this perturbation would allow us to observe alterations in HuR’s binding preferences and interaction dynamics. Mirroring the coCLIP findings, fractionation CLIP of unstressed cells revealed typical enrichment in mRNAs for intronic HuR binding events in the nucleus (75%) and primarily 3’UTR binding events in the cytoplasm (60%) (**Figure 2A**). While coCLIP under stressed conditions maintained intronic preference in the nucleus and 3’UTR preference in the cytoplasm, arsenite stress led to a balanced proportion of intronic and 3’UTR binding in fractionation CLIP (**Figure 2A**). For non-coding HuR targets, fractionation CLIP and coCLIP presented similar proportional distributions under normal conditions (**Figure 2B**). In contrast, we observed a specific increase in cytoplasmic coCLIP under stress conditions for transposable elements (TEs), snRNAs, and snoRNAs. Overall, these findings indicate that arsenite can impact the results derived from fractionation methods, leading to changes in observed HuR binding patterns while emphasizing coCLIP’s consistent performance under these conditions.

**Figure 2:**
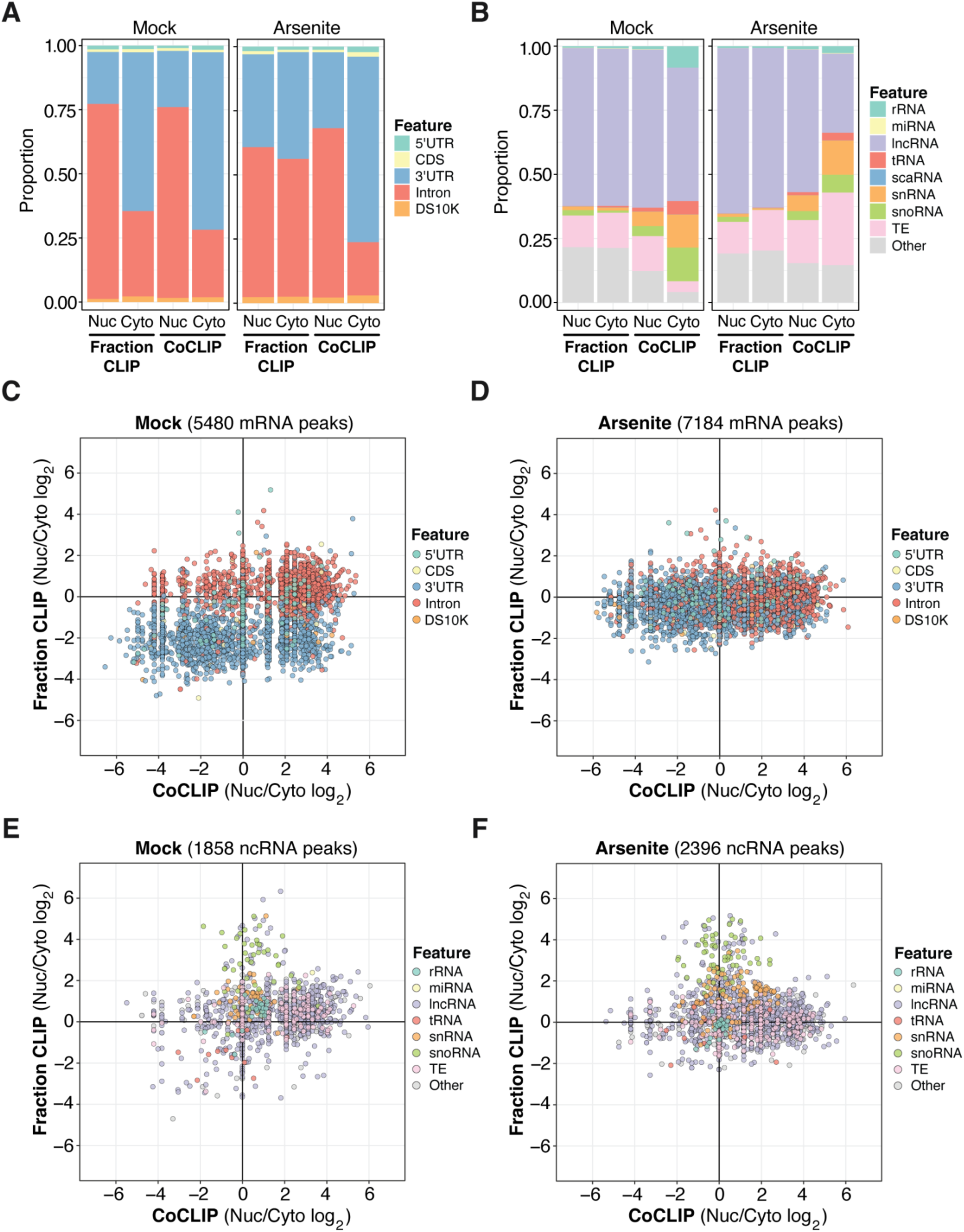
Consistency of HuR binding preferences with coCLIP under arsenite stress. **(A)** HuR peak proportions under mock and arsenite stress conditions for nuclear or cytoplasmic CLIP derived from classical fractionation or in situ labeling with coCLIP. **(B)** HuR peak proportions for ncRNAs as in **(A)**. **(C-D)** Scatterplots comparing nuclear to cytoplasmic ratios of HuR mRNA binding in coCLIP and fractionation CLIP during mock **(C)** and arsenite **(D)** stress conditions. **(E-F)** Similar scatterplots as in **(C-D)** for HuR binding on ncRNAs during mock **(E)** or arsenite **(F)** stress conditions. See also Figure S4.

We next sought to directly compare nuclear and cytoplasmic enrichment for target HuR peaks in fractionation CLIP and coCLIP. Under mock conditions, we identified clear, annotation-dependent enrichment of nuclear introns and cytoplasmic 3’UTR targets (**Figure 2C**), a pattern preserved exclusively in coCLIP during arsenite stress (**Figure 2D**). For noncoding HuR targets, we noticed similar distributions between nuclear and cytoplasmic locations under both conditions; however, coCLIP demonstrated a pronounced nuclear enrichment for lncRNAs, snRNAs, and TEs in comparison to fractionation CLIP (**Figure 2E-F**). We observed a modest nuclear enrichment for snRNAs in coCLIP data that was reduced upon stress, while snoRNAs were enriched in nuclear fractionation CLIP regardless of stress. These results suggest that while fractionation can produce anticipated outcomes under normal conditions, introducing arsenite stress may generate fractionation artifacts. By performing labeling in situ, coCLIP appears to circumvent these distortions.

### HuR exhibits unique binding preferences in stress granules via coCLIP

Our results reveal a consistent alignment between coCLIP and fractionation CLIP in unstressed conditions. However, they also highlight the inherent limitations of fractionation in accessing specific cellular compartments, possibly due to artifacts or other limitations. Many compartments, particularly membrane-less organelles such as SGs,^33,34^ are often transient, condition-dependent, and challenging to purify.^35^ CoCLIP could offer a means to delve into these structures, allowing investigations even in their unstressed states. Previous APEX-based studies of SG proteomes^14^ and transcriptomes^18^ serve as enabling precedents, laying a solid foundation for the deployment of coCLIP to probe specific RBP:RNA interactions and to discern the particular HuR targets residing within SGs with greater precision.

We performed HuR coCLIP using APEX tagged to the SG core protein G3BP1^21^ to study HuR-RNA interactions specific to SGs. Peak enrichment analysis revealed pronounced 3’UTR enrichment in SGs (**Figure 3A**). Interestingly, we observed 3’UTR enrichments in SG in both mock and arsenite treatment conditions, suggesting a basal level of sub-microscopic SG formation with nucleoprotein complexes in the absence of cellular stressors. Moreover, we observed a unique enrichment of TEs within SGs upon arsenite treatment, consistent with the previously observed accumulation of LINE/SINE elements like ALU in SGs (**Figure 3B**).^36^

**Figure 3:**
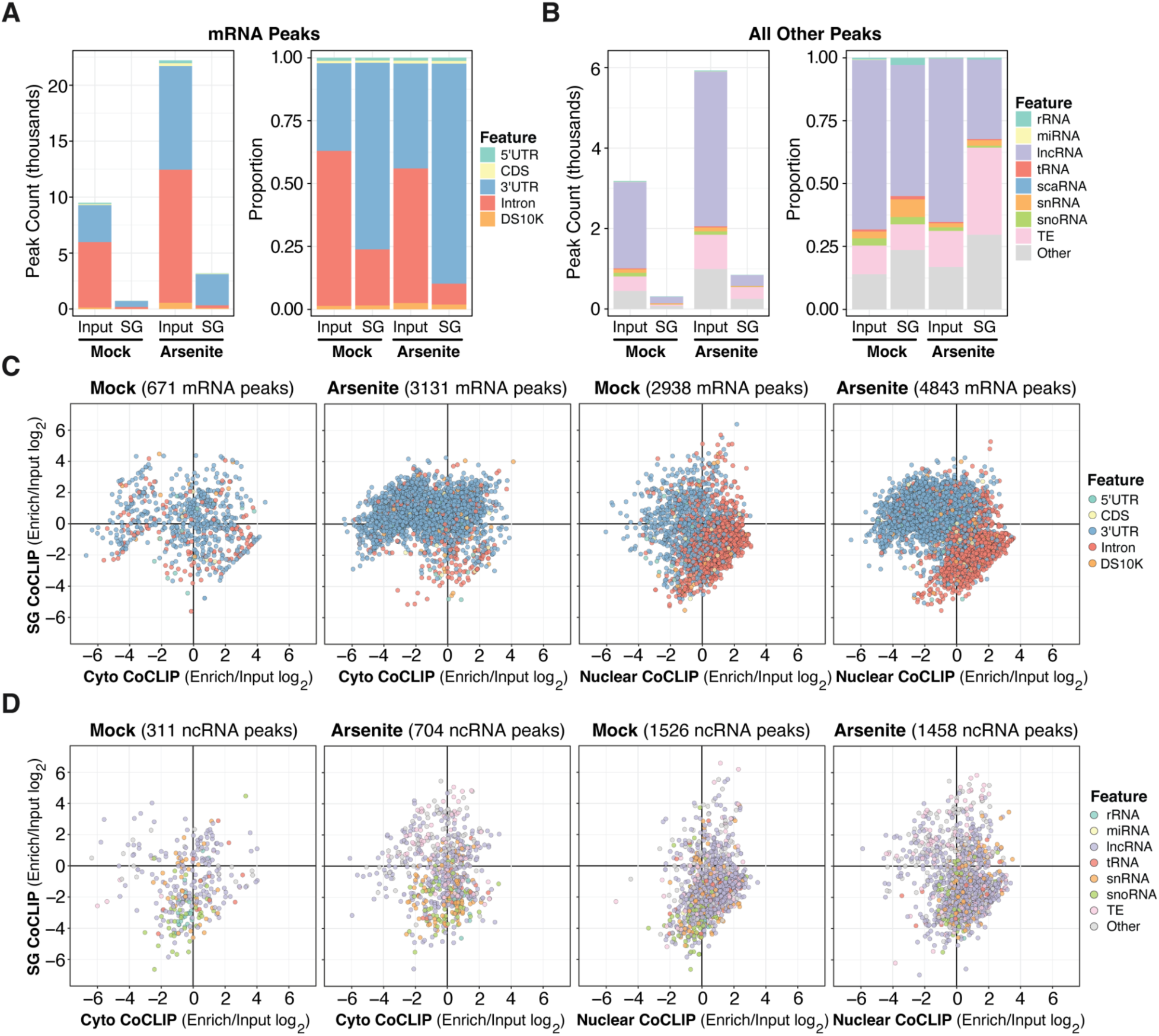
HuR exhibits unique binding preferences within stress granules. **(A)** HuR peak counts (left) and proportions (right) on mRNAs for input CLIP or SG coCLIP in Huh7.5 cells. DS10K: within 10kb downstream of 3’eUTRs. **(B)** HuR peak counts (left) and proportions (right) as in **(A)** for ncRNAs. **(C)** Scatterplots contrasting enriched coCLIP to input CLIP for mRNA binding events, comparing SG HuR to cytoplasmic or nuclear coCLIP under mock and arsenite stress conditions. **(D)** Scatterplots as in **(C)** for ncRNAs.

Next, we analyzed the enrichment of HuR peaks in the nucleus, cytoplasm, and SGs, comparing them to input HuR CLIP under mock and arsenite-induced stress conditions. Under arsenite treatment, HuR in both the cytoplasm and SGs exhibited heightened 3’UTR enrichments relative to input HuR CLIP (**Figure 3C**). However, the HuR specific to SGs displayed more pronounced 3’UTR peak enrichments, which were absent in the cytoplasmic HuR. In both conditions, the nuclear-specific HuR maintained its typical predominance for intronic peaks. Moreover, compared to its nuclear or cytoplasmic counterparts, the SG-specific HuR revealed elevated TE peak binding (**Figure 3D**) without similar enrichments in other non-coding RNAs (e.g., snoRNA, snRNA, rRNAs, and lncRNAs). These results suggest that HuR exhibits localization-specific target enrichment and binding preferences.

### HuR demonstrates localization and condition-specific binding preferences across genomic loci and motifs

To further characterize HuR binding preferences across distinct cellular compartments, we conducted a metagene analysis on five genomic loci: the transcription start site, translation start and stop sites, and splicing donor/acceptor sites (**Figure 4A**). For nuclear HuR coCLIP, we noticed HuR binding enrichment near 5’ and 3’ splice sites with reduced signal in the 3’UTR, a trend consistent between both mock and stress conditions. Cytoplasmic HuR coCLIP showed more robust enrichment for peaks in 3’UTR compared to the nucleus. Intriguingly, in mock conditions, cytoplasmic HuR also demonstrated binding proximal to splice sites, a feature that persisted upon stress, albeit at reduced levels. These results imply that the translocation of HuR from the nucleus to the cytoplasm does not grossly affect normal intronic binding. HuR labeled in stress granules distinctly favored 3’UTRs under both conditions, corroborating our observations in the peak enrichment analysis (**Figure 3A**).

**Figure 4:**
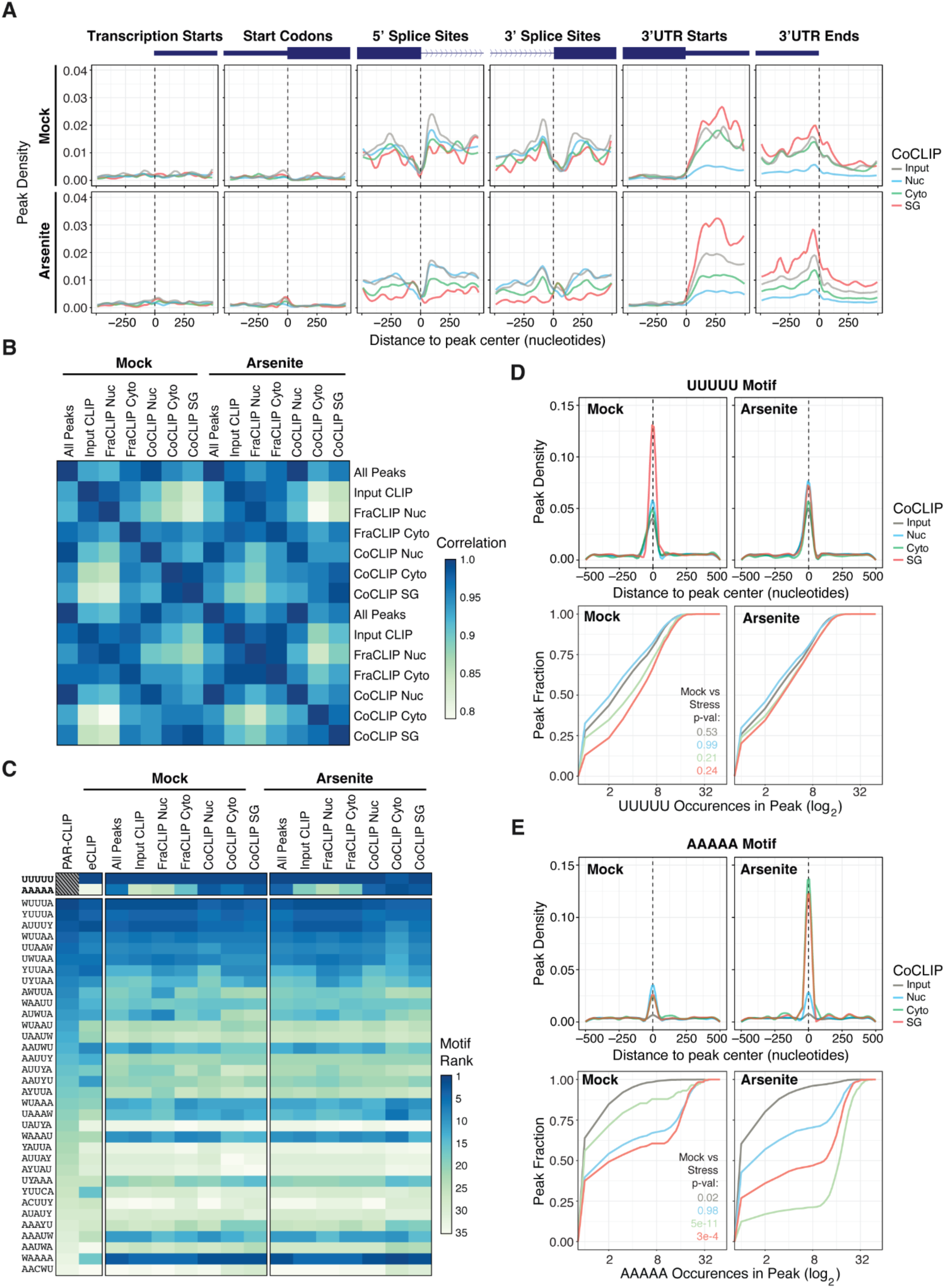
Genomic position and motif preferences for subcellular HuR. **(A)** Metagene profiles for HuR binding for input CLIP or coCLIP under mock or arsenite conditions centered on indicated mRNA features: transcription starts, start codons, 5’ and 3’ splice sites, 3’UTR starts, and 3’UTR ends. **(B)** Correlation matrix for HuR peak binding for all CLIP experiments by type. Frac CLIP: Fractionation CLIP. **(C)** HuR binding motif ranks for all CLIP and coCLIP experiments ordered by PAR-CLIP rankings from ^26^. **(D)** Motif density within HuR peaks (top) and cumulative distribution for frequency of the UUUUU motif (bottom) in input CLIP or coCLIP in mock or stress conditions. **(E)** Motif density within HuR peaks (top) and cumulative distribution for frequency of the AAAAA motif (bottom) as in **(D)**. Indicated P-values calculated by Kolmogorov–Smirnov test.

Depending on localization, differential binding preferences for HuR were also observable at the motif level. We first calculated 5-mer motif densities around peaks (**Table S2**). We utilized 34 target motifs previously characterized for HuR through PAR-CLIP,^26^ and we determined the ranked order of motif enrichments for our datasets. While we observed strong sample correlations between our samples as expected (R^2^ > 0.8, **Figure 4B**), we discerned intriguing differences between samples. Notably, the distinctions between nuclear and cytoplasmic fractionation CLIPs diminished under the influence of arsenite treatment. Conversely, the differences between nuclear, cytoplasmic, and SG CoCLIPs became more pronounced under stress.

The differences observed in the correlation matrix were more apparent when we investigated our samples at the individual motif level (**Figure 4C**). Generally, we observed that the rank order of the enriched motifs from our experiments matched well with what has been reported in the field,^7,23,26^ especially for uridine-rich motifs. Unexpectedly, we detected a pronounced enrichment of adenine-rich motifs compared to published data. While A-rich motifs such as ’WAAAA,’ ’AAAUW,’ ’WAAAU,’ ’UAAAW,’ and ’WUAAA’—previously identified as HuR targets—showed elevated enrichment levels, the ’AAAAA’ motif showed strong enrichment exclusively in the CoCLIP samples.

Analysis of motif density within peaks revealed a strong representation of the ‘UUUUU’ motif across coCLIP samples and conditions (**Figure 4D**), with the most significant binding to the ‘UUUUU’ motif observed in SG HuR under mock conditions. In addition, the cumulative distribution of ‘UUUUU’ appearances within peaks showed that roughly 50% of peaks containing ‘UUUUU’ have four or fewer motifs (including overlaps) under all samples and conditions and exhibited no statistically significant differences.

In contrast, adenine-rich motifs were enriched within HuR peaks in the cytoplasm and SGs under stress conditions (**Figure 4E**). The cumulative distribution of ‘AAAAA’ appearances within cytoplasmic and SG peaks displayed statistically significant shifts between mock and stress conditions (Kolmogorov-Smirnov test, p < 0.01). While 50% of peaks in cytoplasmic HuR contained at most one ‘AAAAA’ motif, the number of ‘AAAAA’ motifs significantly increased upon stress. SG HuR showed a similar shift in which 50% of peaks in the mock condition contained two or fewer ‘AAAAA,’ while upon stress, 50% contained at least eight ‘AAAAA’ motifs. These findings suggest that while HuR binds to its canonical AU-rich elements such as ‘UUUUU’ and ‘AAAAA,’ this binding may not be equivalent. Specifically, our data suggest that HuR binding to adenine-rich motifs is location and condition-specific. The sequence motif context is distinct from uridine-rich motifs, where HuR-bound adenine-rich motifs are considerably longer than uridine-rich motifs upon stress.

### Gene level analysis of subcellular HuR targeting reveals distinct gene set enrichments

Expanding our analysis beyond peak-level enrichment, we honed in on the interaction between HuR and RNA at the gene level. Our primary objective was to eliminate the possibility that the distinct HuR binding patterns we observed in specific locations were artifacts of transcriptomic alterations among different APEX-cell lines or between mock and arsenite treatment conditions. To this end, we conducted RNA-Seq and differential gene expression analysis on the cell lines utilized for CoCLIP experiments, finding no statistically significant disparities between cell lines or our two conditions (**Figure S5**). Target gene enrichment comparisons between CoCLIP and RNASeq of the respective APEX cell lines demonstrated that arsenite treatment was not a source of significant transcriptional difference (**Figure 5A**). Thus, the change to HuR targeting is likely occurring at the level of relocalization. Moreover, HuR target genes exclusively enriched in SG showed longer overall transcript lengths than targets enriched in the nucleus or cytoplasm (**Figure 5B**), consistent with previous observations.^35^

**Figure 5:**
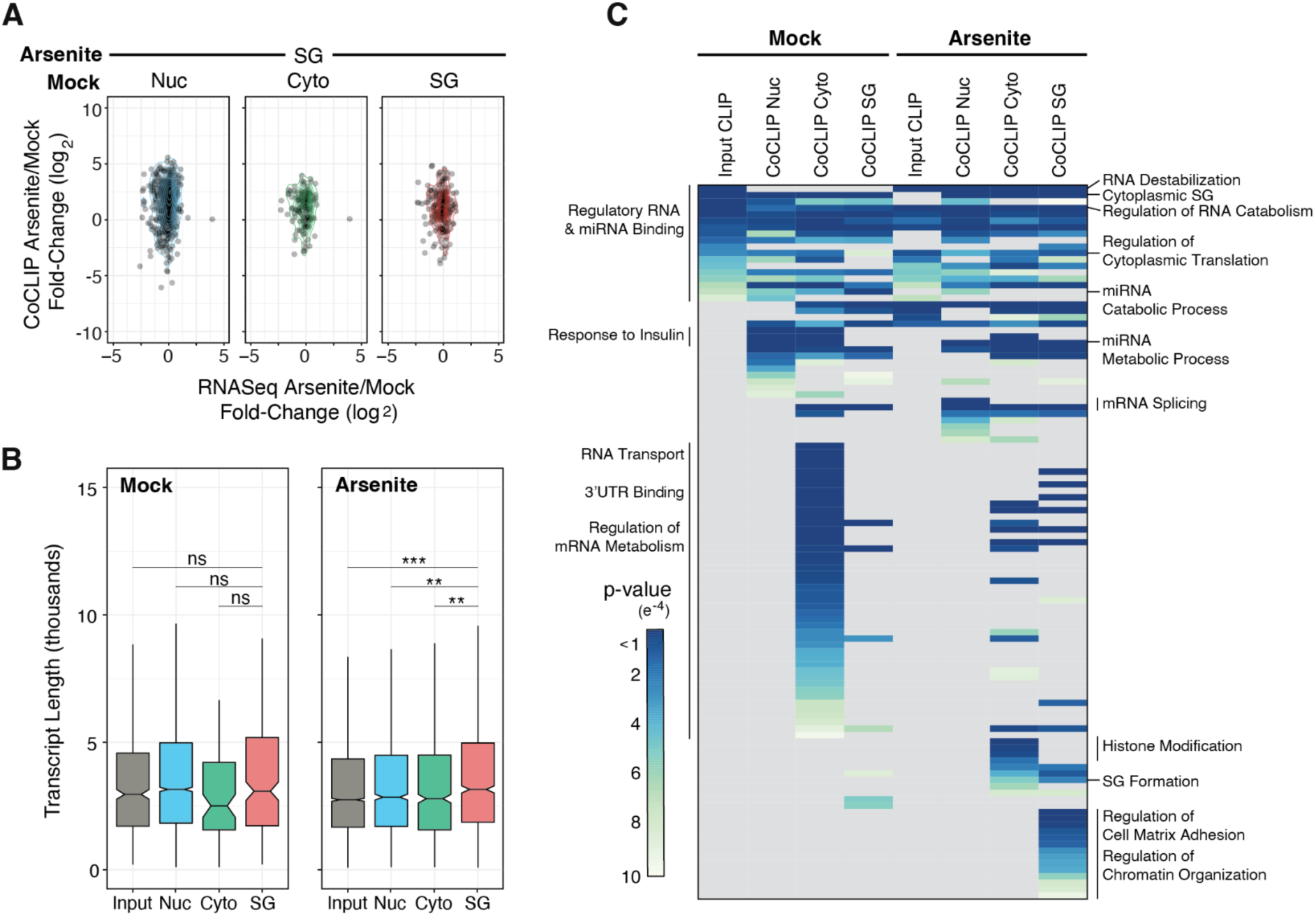
Gene level features of subcellular HuR targeting from coCLIP. **(A)** Stress versus mock comparisons of SG coCLIP data with respective mock conditions compared to matched gene-level RNAseq data. **(B)** Transcript length boxplots for input or coCLIP-derived HuR targets. **(C)** Gene ontology analysis of HuR binding sites from input CLIP or compartment-specific coCLIP under mock or arsenite stress conditions. Notable gene ontology terms are highlighted. ***P<0.001, **P<0.01, ns P>0.05, two-sided Mann-Whitney U-test. See also Figure S5.

We next performed gene ontology enrichment analysis on the HuR target genes identified for different cellular compartments (**Figure 5C**) to uncover potential associations attributed to HuR targets within different cellular environments. All target gene ontologies identified in input mock HuR CLIP were identified in at least one coCLIP dataset, suggesting a level of conserved targeting. However, unique clusters of target genes became apparent under stress and cellular localization. For instance, genes associated with SG assembly were prominently enriched in SG CoCLIP and, to a lesser extent, in cytoplasmic CoCLIP under stress. Still, they were not present in nuclear CoCLIP under stress or in any CoCLIPs under mock conditions (**Figure 5C**). These results shed light on the dynamics of HuR targeting and hint at a structured network where distinct subcellular RNA targets are functionally interconnected.

### De novo HuR binding events and HuR flow across compartments

Our gene set enrichment analysis prompted us to investigate how the HuR translocation from one compartment to another affects its target gene repertoire. As our analysis is far from exhaustive by focusing on three compartments, we also wanted to account for HuR targeting events that appeared to be “de novo” upon stress. These targeting events could be due to HuR:RNA translocation from other compartments as well as new binding events from HuR not bound to RNA. By dividing the identified HuR target genes into subgroups exclusive to each compartment and condition, we tracked target genes shared between mock and stress conditions in the same compartment or genes that were in one compartment in the mock condition but appeared in a different compartment upon stress. To our surprise, almost all (748/763 or 98%) HuR target genes uniquely enriched in nuclear coCLIP under stress conditions were de novo targets not associated with HuR in any tested compartment in the mock condition (**Figure 6A**).

**Figure 6:**
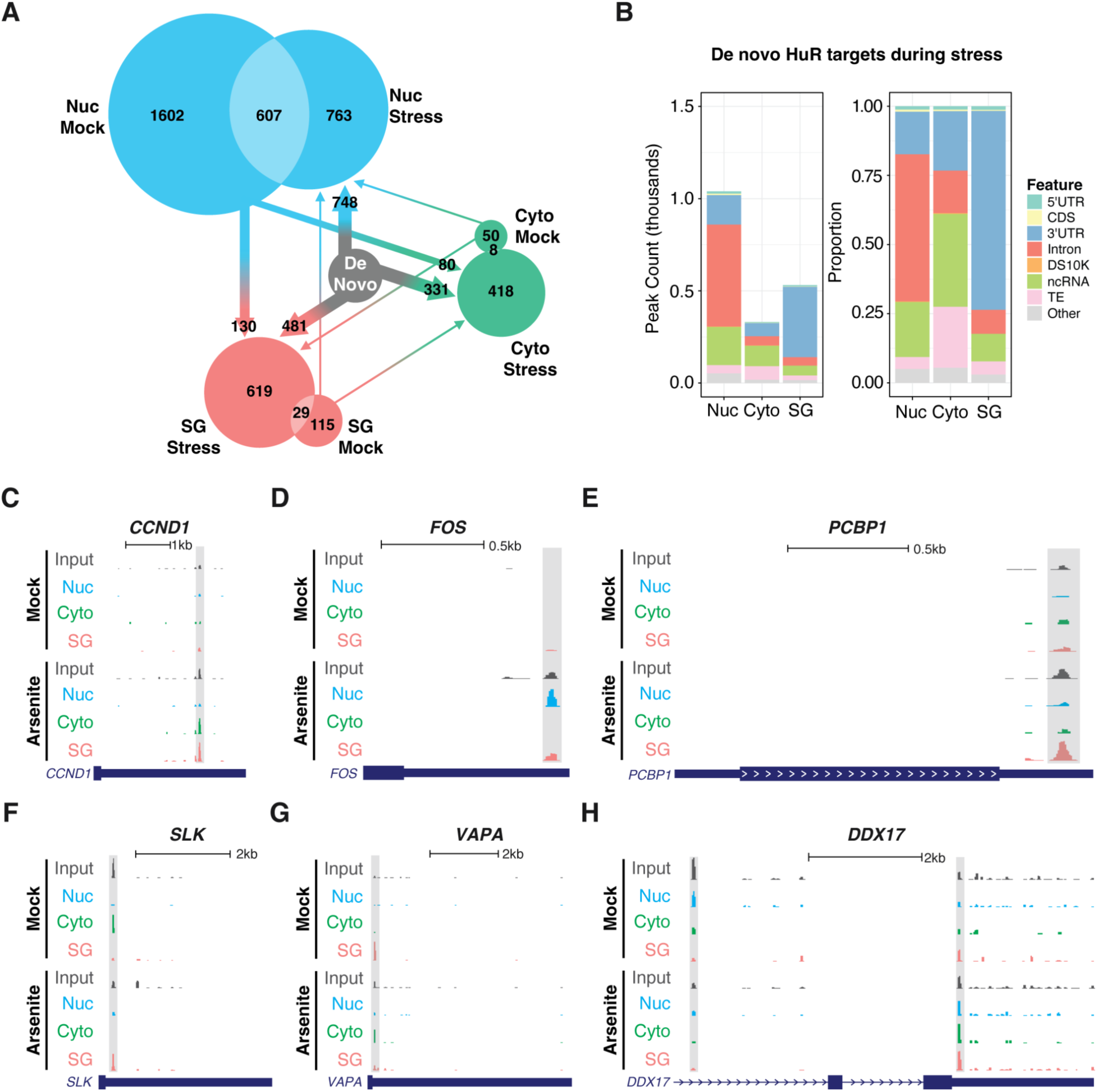
De novo HuR binding events and HuR flow across compartments. **(A)** Graphical illustration of common and unique HuR targeting events from nuclear, cytoplasmic and SG coCLIP under mock and arsenite stress conditions. Arrows indicate the putative source of binding events from the mock condition. **(B)** De novo HuR binding event counts (left) and proportions (right) for nuclear, cytoplasmic and SG coCLIP datasets. DS10K: within 10kb downstream of 3’UTRs. **(C-H)** Gene browser tracks of normalized HuR binding for input CLIP and coCLIP data on the **(C)** *CCND1* 3’UTR, **(D)** *FOS* 3’UTR, **(E)** *PCBP1*, **(F)** *SLK* 3’UTR, **(G)** *VAPA* 3’UTR, and **(H)** *DDX17* introns and 3’UTR. Significant peaks are highlighted in light gray.

Similarly, approximately 80% of HuR target genes unique to SGs and the cytoplasm during stress were de novo targets. Furthermore, nuclear HuR target genes uniquely enriched in mock conditions were also targets in the cytoplasm and SG under stress conditions (80 and 130 genes, respectively). In contrast, only a small number of cytoplasmic and SG HuR target genes under mock conditions were observed in different compartments under stress. Within each compartment, 20% of nuclear target genes were consistent between mock and stress conditions, whereas less than 5% of the target genes in the cytoplasm and SGs were shared across conditions.

The de novo target genes for each compartment under stress conditions showed peak distributions (**Figure 6B**) that were characteristic of each compartment (**Figure 2A**, **Figure 3A, B**). Notably, the enhanced TE peak binding observed in SG HuR under stress was absent among the de novo target genes. These results imply that HuR association with TEs is distributed evenly across the cellular compartments we’ve tested under mock conditions.

To further illustrate these variations in HuR binding, we examined individual gene tracks across different compartments and stress conditions. For example, HuR showed minimal binding on mRNA for the cell cycle regulator cyclin D1 under mock conditions (**Figure 6C**). However, we observed strong HuR binding in the 3’UTR of *CCND1* mRNA in the input HuR CLIP upon stress induction. Through coCLIP, we discerned that this HuR binding predominantly occurred in the cytoplasm and within SGs, with no detectable binding in the nucleus. Likewise, the proto-oncogene Fos demonstrated no interaction with HuR across all compartments under mock conditions; however, under stress conditions, HuR binding to the 3’UTR of the *FOS* transcript was evident, specifically in the nucleus and SGs (**Figure 6D**). We also observed similar HuR binding patterns on the PCBP1 transcript, which encodes a multifunctional RBP implicated in alternative splicing, immune regulation, and cancer (**Figure 6E**), where increased association with HuR in the 3’UTR under stress conditions was primarily due to increased HuR binding in SG.

Furthermore, coCLIP data were sensitive enough to discern instances where HuR binds to identical locations on a transcript during both mock and stress conditions but in different cellular compartments. For instance, the mRNA for the protein kinase *SLK* exhibited a robust association with HuR in the 3’UTR of its gene in the cytoplasm under mock conditions (**Figure 6F**). However, with arsenite-induced stress, while the HuR binding site remained constant, the interaction location transitioned from the cytoplasm to SG. Conversely, the *VAPA* transcript, encoding the endoplasmic reticulum membrane component VAP-A, demonstrated pronounced 3’UTR binding with HuR in SG under mock conditions (**Figure 6G**). Following arsenite treatment, this same binding interaction predominantly occurs in the cytoplasm. Lastly, our data could delineate even more intricate alterations in HuR binding that occur across different loci on a transcript, varying between subcellular compartments and conditions. For example, the *DDX17* mRNA displays robust intronic HuR binding in the nucleus under mock conditions. However, upon the induction of stress, this intronic binding is reduced, but there is a corresponding increase in the 3’UTR of the gene in the cytoplasm and SGs (**Figure 6H**). Collectively, our findings demonstrate that coCLIP is a robust and sensitive innovative method for studying RNA-RBP interactions with subcellular resolution.

## Discussion

Here, we report a novel method, coCLIP, that allows the investigation of RNA-RBP interactions at subcellular compartment resolution by combining CLIP with APEX2 proximity labeling. We used HuR, a ubiquitous RBP with known nuclear and cytoplasmic specific functions and binding profiles, as a benchmark for the utility of coCLIP. Our results show robust recapitulation of HuR binding patterns under normal cell growth conditions, with enriched binding to intronic regions in the nucleus and to 3’UTRs in the cytoplasm.

We observed a notable disparity between the results obtained from fractionation and coCLIP, particularly under arsenite-induced stress conditions. While fractionation methods demonstrated reliable results under normal (mock) conditions, this resolution was compromised under arsenite stress, displaying an equal proportion of intronic and 3’UTR binding, potentially indicative of alterations in binding profiles. Conversely, coCLIP, with its in situ labeling approach, maintained consistent results under both mock and stress conditions, preserving the intronic and 3’UTR preferences. We interpret these results to suggest that coCLIP is more resilient to changes brought about by stressors like arsenite and, as such, offers a more accurate picture of RNA-protein interactions within subcellular compartments.

Our coCLIP experiments with HuR highlight the intricacies of binding patterns influenced by the subcellular location of HuR and the cellular context. We observed a discernible shift in target sets at the gene level between mock and arsenite stress conditions. Yet, at a more granular level, HuR’s binding preferences exhibit adaptability within individual genes, moving to different but specific loci under varying conditions. For instance, the translational arrest induced by arsenite and other integrated stress response (ISR) agonists is thought to result in free mRNA that acts as a nucleation signal for G3BP and other mRNA binding RBPs.^33,37–40^ We notably did not observe HuR binding to coding regions during stress but did observe an increase in 3’UTR target binding, both in terms of the number of targets and intensity at specific target loci. These results suggest that client proteins such as HuR may remain stably bound to their targets as they are redirected to SGs from other compartments, and perhaps that true “de novo” HuR binding remains directed to specific loci within SGs. As the landscape of RNA-protein interactions and the resultant regulatory behaviors are shaped by a combination of factors, including the availability co-binding RBPs and target RNAs, our findings underscore the need for detailed studies to unravel the specific cues guiding RBP target selection and localization, especially under stress conditions.

Based on these observations, coCLIP opens avenues for addressing intriguing biological questions centered on the regulatory dynamics of RNA-RBP interactions. For instance, what factors influence HuR’s choice of one genomic locus over another, and how does subcellular localization affect this choice? Exploring these questions from a protein-centric viewpoint could offer more insights, examining how post-translational modifications of HuR or alterations in the availability of proteins that bind cooperatively or competitively to similar motifs in the cytoplasm and stress granules affect these interactions. Additionally, it is worth exploring whether this observed binding pattern shift is exclusive to HuR or represents a shared behavior across other nucleocytoplasmic shuttling RBPs^41^ in additional stress contests,^42^ which could further enrich our understanding of RBP:RNA interaction networks.

Another important question is the relationships between changes in HuR binding and downstream functional roles within different cellular compartments. In addition to binding target gene sets that are shared across compartments and conditions, we observe that HuR binds distinct sets of genes in a location and context-specific manner. These results allow us to envision a framework in which the localization of HuR, in tandem with changes in local RNAs or protein concentration, influences RBP binding and, ultimately, downstream regulatory outcomes. This perspective corroborates regulatory models, such as RNA regulons^43^ and matchmaking RBPs,^44^ that describe how an RNA-protein complexes may coordinate the regulation of functionally related mRNA transcripts. CoCLIP presents a means to explore components of these intricate regulatory models at a subcellular level, potentially clarifying the mechanisms governing coordinated RBP:RNA interaction networks.

### Limitations of This Study

While coCLIP advances our understanding of RNA-RBP interactions, its current design interrogates one protein within a designated compartment at a time. This specificity means that comprehensive insights, such as HuR target RNA interactions with additional RBPs in other compartments, remain to be explored. Further, in line with other proximity-based techniques, coCLIP may capture RBPs initially labeled in one compartment but subsequently trafficked to another. For instance, we observed nuclear non-coding RNAs such snoRNAs and snRNAs in cytoplasmic HuR CoCLIP. This is likely be due to cytoplasmically labeled HuR moving to the nucleus as UV crosslinking took place. While such instances appear limited in our data, they emphasize the importance of understanding RBP dynamics across compartments.

Encouragingly, advances in dual labeling approaches^45^ hold promise in enhancing the accuracy of future studies.

## Acknowledgements

We are grateful to Charlie Rice for his unwavering support, mentorship, and guidance. We thank Eckhard Jankowsky, Maria Hatzoglou, Kristian Baker, Caryn Hale, Ezgi Hacisuleyman, Erin Conlon and Robert Darnell for technical advice and helpful feedback. We further thank Frank Tedeschi and Thomas J. Sweet for feedback and a critical reading of this manuscript. We gratefully acknowledge the Genomics Core Facility at the Genetics and Genome Sciences Department at Case Western Reserve University School of Medicine and the Genomics Resource Center at The Rockefeller University for their services. We further thank the members of the Rice and Luna labs for advice and support. This work was supported in part by the NIH under the award T32 GM007250 (S.Y.), funding from the Case Comprehensive Cancer Center P30 CA043703 (J.M.L), the American Cancer Society IRG-16-186-21 (J.M.L), as well as startup funds from the Department of Biochemistry at the Case Western Reserve University School of Medicine.

## Author Contributions

Conceptualization, J.M.L; Methodology, S.Y, K.R-G, and J.M.L.; Investigation, S.Y, S.S.S, and J.M.L; Writing – Original Draft, S.Y and J.M.L..; Writing – Review & Editing, S.Y and J.M.L..; Funding Acquisition, J.M.L.; Supervision, J.M.L.

## Declaration of Interests

The authors declare no competing interests.

## Materials and Methods

### Plasmid DNA construction

To generate stable cell lines expressing APEX2 constructs, we cloned FLAG-APEX2 (Addgene, #92158) into enhanced Piggybac (ePB) doxycycline-inducible expression vectors.^46^ Overlap PCR constructs encoding FLAG-APEX2 fusions with mKate2 fluorescent protein (Evrogen, cat. #FP181) were cloned between BamHI and NotI sites in ePB vectors as follows: for cytoplasmic APEX, the nuclear export signal (LALKLAGLDI) from PKI^47^ was appended to the N-terminus of FLAG-APEX2-mKate2; for nuclear APEX, the nuclear localization signal (PKKKRKV) from SV40 LT^48^ was appended to the N-terminus of FLAG-APEX2-mKate2; for ER-localized APEX2, we cloned DNA encoding amino acids 1-60 from human Sec63^49^ from cDNA and appended this sequence at the N-terminus of FLAG-APEX2-mKate2; for stress granule APEX2, we fused G3BP1 containing the S149E mutation^50,51^ to the C-terminus of FLAG-APEX2-mKate2. All primers to generate these PCR amplicons are listed in table S3.

### Cell culture and stable cell line generation

Huh-7.5 cells (*H. sapiens*; sex: male),^52^ were maintained at 37 °C and 5% CO_2_ in Dulbecco’s Modified Eagle Medium (DMEM, Fisher Scientific, cat. #11995065) supplemented with 0.1 mM nonessential amino acids (NEAA, Fisher Scientific, cat. #11140076) and 5% hyclone fetal bovine serum (FBS, HyClone Laboratories, Lot. #AUJ35777). All cell lines have tested negative for contamination with mycoplasma.

Stable cell lines were generated by transfecting 1μg of ePB plasmid encoding localized APEX2 with 1μg of pTransposase plasmid^46^ using Lipofectamine 2000 (Invitrogen, cat. #11668027). Cells underwent selection with 2μg/ml Puromycin (Sigma, cat. #P8833-25MG) to eliminate non-transfected cells. Doxycycline (Sigma, cat. #D9891-1G) was used at 1μg/ml to induce APEX2 expression.

### APEX labeling and crosslinking

Cells were plated onto 10 or 15 cm dishes, and expression was induced for 2 days with 1μg/ml of doxycycline. At 45 minutes prior to APEX labeling, sodium arsenite was added to the media to a final concentration of 0.5mM and plates were placed back into the incubator. At 30 minutes prior to APEX labeling, biotin-tyramide (Iris Biotech, cat. #LS-3500.1000) was added to a final concentration of 500μM and plates were placed back into the incubator. APEX labeling was initiated at room temperature by adding H_2_O_2_ at a 1mM final concentration to each dish. Cells were gently agitated for 1 minute, and quenched twice in ice-cold quenching buffer (5mM Trolox and 10mM sodium ascorbate in DPBS). The quenching buffer was replaced with ice-cold PBS and cells were immediately irradiated under 254nm UV light on ice, once for 400 mJ/cm^2^ and once again for 200 mJ/cm^2^ using a Spectrolinker XL-1500 (Spectronics Corporation). PBS was then replaced with quenching buffer. Cells were scraped into tubes and pellets were stored at −80°C for coCLIP.

### CLIP and CoCLIP

CLIP from crosslinked Huh7.5 cell pellets was performed generally following previous work^53,54^ with modifications for CoCLIP.

#### Bead preparation

Two sets of beads were prepared, one set for HuR antibody-based pull-down and a second for streptavidin pulldown. For HuR antibody pull downs, protein G Dynabeads (Invitrogen, cat. #10004D) were washed 3x with and resuspended in antibody binding (AB) buffer (AB: PBS, 0.02% Tween-20). Per sample, 50μL of beads was used. Beads were incubated with 3μg of HuR 3A2 antibody (Santa Cruz, cat. #sc-5261) per 50μL beads for 30 min at room temperature or overnight at 4°C. Prior to IP, beads were washed in 1x PXL lysis buffer (1x PXL: 1X PBS tissue culture grade without magnesium or calcium, 0.1% SDS, 0.5% Sodium-deoxycholate, 0.5% NP-40, with protease inhibitors). For streptavidin pull-down, we equilibrated 50μL of MyOne T1 beads (Invitrogen, cat. #65601) per sample in 1x PXL containing no SDS.

#### HuR IP and on-bead enzymatic steps

Lysates from crosslinked cells were prepared by adding 1ml of 1x PXL lysis buffer + quenchers + protease inhibitors (Roche, cat. #11873580001) and triturating to disrupt cells. Lysates were treated with 10μL DNase (RQ1, Promega) and placed on ice for 10 minutes. Lysates were then treated with RNase I (Thermofisher, cat. #EN0602), first diluted to the indicated concentration by volume (e.g., 1:100 for “over digestion or OD”, or 1:5,000 for ‘‘Low RNase”) in 1x PXL and then added at 10μl per mL of lysate. Lysates underwent thermomixing for 5 minutes at 37°C at 1100rpm before being spun at 4°C on max speed of a table-top microcentrifuge for 10 minutes. All subsequent steps were done on ice or at 4°C unless otherwise indicated. Supernatants, along with any lipid layer, were harvested and mixed with PXL-equilibrated HuR antibody-bound beads for immunoprecipitation. Samples were nutated with beads at 4°C for 2-4 hours. Beads were washed sequentially twice each with ice-cold 1x PXL, 5x PXL (same as 1x but using 5x PBS), and 1x PNK buffer (50mM Tris-HCl, pH 7.5, 10mM MgCl2, 0.5% NP-40).

To prepare RNA 3’ ends for linker ligation, IPs were treated with alkaline phosphatase. Beads were resuspended in 40μL containing 1x dephosphorylation buffer, 3U of CIAP (Roche), RNasin inhibitor (Promega), and thermomixed for 20 minutes at 37°C, shaking at 1100rpm for 15 s every 2 min. Samples were washed as above sequentially in ice-cold 1x PNK, 1x PNK plus 20mM EGTA, and twice with 1x PNK.

Radiolabeled linkers were prepared with a poly-nucleotide kinase (PNK) reaction consisting of 100pmol of a 3’ inverted ddT blocked L32 RNA linker, 0.5μL of 32P-gamma-ATP (Revvity, cat. #NEG035C005MC), 0.5µl 1x T4 PNK buffer, 0.25μL T4 PNK and 0.3μl RNasin inhibitor in a total volume of 5µl per sample. Radiolabeled linker was usually prepared for 10 ligation reactions in a volume of 50µl. Linkers were incubated for 20 min at 37°C, after which 2µl of 10mM ATP was added, and the reaction was incubated at 37°C for an additional 5 minutes. Linker was purified by passing through a G-25 column (GE Healthcare, cat. # 27-5325-01) and stored at −20°C until use.

Linker ligation at 3’ ends was set up per sample after washes following alkaline phosphatase treatment by preparing a T4 RNA ligase 1 (NEB, cat. #M0204S) reaction in 40μL following the manufacturers’ instructions with 100pmol of radiolabeled L32 RNA linker. Samples were incubated overnight at 16°C, shaking at 1100rpm for 15 s every 4 min. The next day, beads were washed twice each with 1xPXL and 5x PXL, before being equilibrated in 1x PXL.

#### CoCLIP streptavidin pulldown and autoradiography

Beads were then subdivided into two fractions: For total HuR CLIP, 20% of the beads were set aside on ice. To the remaining 80% of beads, samples were resuspended in 1x PXL containing 1% SDS and HuR was eluted via incubation at 70°C for 10 minutes. Eluted HuR supernatant was added to an equal volume of MyOne T1 beads equilibrated in 1x PXL containing no SDS, for a final SDS concentration of 0.5%. Samples were incubated at 4°C nutating for 2 hours. All eads were subsequently washed twice each with 1x PXL, 5x PXL, and 1x PNK. Protein was eluted off the beads by incubating with 30µl of 1X LDS loading buffer (Invitrogen) without reducing agent for 10 min at 70°C, shaking at 1100rpm. Biotin was added to a final concentration of 1mM for all streptavidin IP samples prior to heat elution. Supernatants were run on Novex NuPAGE 8% Bis-Tris cels (Invitrogen, cat. #WG1001BOX) in SDS-MOPS buffer at 4°C. Radiolabeled protein RNA complexes were transferred to BA85 nitrocellulose (Cytiva Amersham, cat. #45-004-007). After transfer, the membrane was rinsed with RNase-free PBS, and exposed to Biomax MR film (Kodak) at 70°C typically from 3 hr to up to 3 days. Alternatively, membranes were placed onto phosphor screens (GE cat. # BAS-IP MS 2025), and imaged with a Typhoon Scanner (GE Amersham).

Nitrocellulose membranes were aligned with the exposed film and regions of the membrane from low RNase IP lanes were excised corresponding to signal intensities for HuR:RNA complexes between 45 and 75kDa. RNA was liberated from membrane fragments using 200µL of a 4mg/ml proteinase K solution (Roche, cat. #3115828001) diluted in PK buffer (100mM Tris-HCl, pH 7.5, 50mM NaCl, 1mM EDTA, 0.2% SDS) and incubated for 60 min at 50C, shaking at 1100rpm for 15 s every 2 min. RNA fragments underwent acid phenol:chloroform extraction using and were precipitated overnight at −80°C. RNA was pelleted by spinning at max speed (> 13,000rpm) in a table top centrifuge at 4°C, and washed twice with 75% ethanol. Following the drying of the RNA at the bench, the pellet was dissolved in 8µL RNase-free water.

#### HuR footprint library generation and sequencing

CLIP footprints were reverse-transcribed using the Br-dU incorporation and bead-capture strategy described previously.^55,56^ Indexed reverse transcription (RT) primers were used (listed in Table S3), allowing multiplexing of up to 8 or 22 samples per Miseq or Nextseq 500 run, respectively. cDNA was circularized with CircLigase (Epicentre), pooled and then amplified with PCR primers with Illumina sequencing adapters as desribed previously ^55,56^. Amplification was tracked with SYBR green (Life Technologies) on the Quantstudio real-time PCR machine (Thermo), and reactions were stopped once signal reached 100k relative fluorescence units (r.f.u.). Products were purified with Ampure XP beads (Beckman) and quantified via Qubit assay (ThermoFisher) and/or Tapestation system (Agilent). Multiplexed samples were run on the Illumina Miseq or NExtSeq Mid-Output with 75 base pair single-end reads.

### RNAseq library preparation

RNA sequencing libraries were prepared for the stable cell lines expressing localized APEX2 in mock and arsenite stress conditions. Cellular stress was induced by treating cells with 0.5mM sodium arsenite for 45 minutes. RNA was isolated with trizol reagent and stored at −80C until use. RNA seq libraries were constructed from total RNA using the NEBNext Ultra II Directional RNAseq kit (NEB) with ribosomal RNA depletion, and sequenced on a NextSeq 500 Illumina Sequencer.

### Cell fractionation

Crosslinked cell pellets were fractionated using the REAP method^31^ with modifications. Briefly, freshly pelleted and crosslinked cells were suspended in 500μl of ice-cold REAP buffer (0.1% NP-40, in DPBS with complete protease inhibitors (Roche, cat. #11873580001), triturated 15 times and incubated on ice for 10 minutes. The lysate was then clarified in a chilled microcentrifuge to create a cytoplasmic fraction and nuclear pellet. The ∼500μl cytoplasmic fraction was saved in a separate tube into which 500μl of 1X PXL was added and subsequently used for western analysis or CLIP. The nuclear pellet was washed twice with REAP buffer, and resuspended in 500μl REAP + 500μl 1X PXL before being used for western analysis or CLIP as above.

### Western blots

Cell lysates were prepared in 1x PXL lysis buffer supplemented with complete protease inhibitors (Roche, cat. #11873580001). All lysates were prepared from equal cell numbers per condition. Protein concentrations were determined by Bradford assay (Biorad), and 20μg total protein per sample was run on NuPAGE gels (Life Technologies) and transferred to fluorescence-compatible nitrocellulose membranes (Millipore). Membranes were blocked in Odyssey PBS-based buffer (LI-COR) for 1 hr, then primary antibodies were added for an overnight incubation at 4°C. Antibodies used for western blotting were: HuR 3A2 (Santa Cruz, cat. #sc-5261, 1:1000), TIA1 (Proteintech, cat. #12133-2-AP, 1:500), GAPDH (Cell Signallng, cat. #5174S, 1:1000), ADAR1 D-8 (Santa Cruz, cat. #sc-271854, 1:1000), H3 (Cell Signallng, cat. #9715, 1:1000) and Beta-Actin (Sigma, cat. #A5441, 1:5000). After 3 washes in 1X PBS/0.05% Tween-20, membranes were incubated with fluorescent secondary antibodies (LI-COR, 1:25,000) for 1 hr at room temperature. AlexaFluor 680 conjugated streptavidin (ThermoFisher, cat. # S21378) was used during secondary antibody incubation at 1:10000 dilution to mark biotinylated proteins. Membranes were washed 3 times in 1X PBS/0.05% Tween-20, rinsed in 1X PBS, and visualized on the Odyssey Imaging system (LI-COR).

### Immunofluorescence

Cells were plated either on glass coverslips or black-walled clear bottom 96-well plates (Corning, Cat# 3904) that were coated with poly-L-lysine. Upon harvest, cells were fixed with 4% paraformaldehyde (PFA) for 10 minutes. PFA was then removed, and cells were stored at 4°C in PBS containing 1% FBS until processing. Cells were washed in PBS containing 0.1% Tween-20 (PBST), permeabilized with PBS containing 0.1% Triton X-100 for 10 min at room temperature, and blocked for 1h at room temperature with a blocking solution of 5% BSA in PBST. Cells were stained with primary antibodies overnight at 4°C. Antibodies used for immunofluorescence were: HuR 3A2 (Santa Cruz, cat. #sc-5261, 1:1000), FLAG M2 (Sigma, cat.: F1804). After primary antibody incubation, cells were washed and stained with secondary antibodies donkey anti-rabbit AlexaFluor 594 (ThermoFisher, cat. #A-21207, RRID:AB_141637) at 1:2,000, donkey anti-mouse AlexaFluor 594 (Abcam, cat. #ab150108, RRID:AB_2732073) at 1:2,000. AlexaFluor 488 conjugated streptavidin (ThermoFisher, cat. # S11223) was used during secondary antibody incubation at 1:5000 dilution to stain biotinylated proteins. Nuclei were counterstained with DAPI (ThermoFisher Scientific cat. #D1306, RRID:AB_2629482) at 1 µg/ml, for 5 minutes prior to imaging. Fluorescent images were obtained on a Keyence BZ-X710 microscope or on a Cytation 7 (Agilent).

### Bioinformatics

Detailed information for all bioinformatic processing with BITs and CLIPittyCLIP in this paper, including relevant shell and R scripts, can be found in the accompanying GitHub repositories related to this manuscript: https://github.com/LunaRNALab/CLIPittyClip and https://github.com/LunaRNALab/BITs.

#### FASTQ processing to finished peak matrix

In order to analyze all our data in a systematic manner, we created a custom single command line tool, *CLIPittyClip*, to process sequencing reads into peaks (**Figure S3**). Briefly, FASTQ files were preprocessed using the fastx_toolkit (http://hannonlab.cshl.edu/fastx_toolkit/) and the CLIP Tool Kit (CTK).^57^ We first collapsed identical reads to reduce redundancy and improve computational efficiency. Unique molecular identifiers (UMIs) were then stripped from each read and appended to read names after which reads underwent demultiplexing to segregate reads based on their barcode sequences. After demultiplexing, barcode and adapter sequences were trimmed to ensure that only high-quality regions of the reads were retained. Reads were at minimum 16nt long for downstream analysis.

Post-preprocessing, the reads were mapped to the human Hg38 reference genome using Bowtie2^58^ allowing zero mismatches. The alignment results, in SAM format, were converted to indexed BAM format using Samtools.^59^ BEDTools^60^ was then used to convert the BAM files into BED format as well as to generate genome-wide coverage files.

Peak calling was performed using HOMER^61^ starting from a tag directory that contained all mapped reads in BED format. Homer peak calling was performed with the following parameters: -style factor -L 2 -localSize 10000 -strand separate -minDist 50 -size 20 -fragLength 25. Identified peaks were further processed to create a standardized BED file: peaks were sorted based on their genomic coordinates and filtered to retain only those on standard chromosomes (1-22, X, Y, and M). Read coverage on peaks across all samples was then calculated using BEDTools. Coverage for each sample was then assembled into a peak matrix for annotation, peak enrichment analysis, and plotting.

#### Peak annotation and prioritization

Peaks were annotated by extracting gene features from the Ensembl release 110 gtf file from GRCh38, as well as RepeatMasked regions (http://www.repeatmasker.org/) from the UCSC table browser.^62^ We first extracted protein-coding genes by gene biotype, and non-coding RNAs (miRNA, lncRNA, rRNA, snoRNA, scaRNA, snRNA, miscRNA) by transcript biotypes. Exons, coding DNA sequence (CDS) exons, introns, 3’ untranslated regions (3’UTRs), and 5’ untranslated regions (5’UTRs) were also extracted. We further calculated a “downstream 10kb” region to account for annotated 3’ ends of transcripts from all 3’UTR endpoints. From RepeatMasker regions we extracted LINE and SINE elements, tRNAs, endogenous retroviral long-terminal repeats (LTRs), low-complexity regions, and satellite repeats. CLIP peaks were then intersected with all the above annotations for overlap. We prioritized peak annotations in the following order: 3’UTR, 5’UTR, CDS, miRNA, lncRNA, rRNA, snoRNA, scaRNA, snRNA, miscRNA, tRNA, TE, Other, CDS_Retained_intron, ncRNA_Retained_intron, intron, downstream 10kb, deep intergenic. TE consisted of the transposable elements LINEs and SINEs, and all other repeat masked regions were summed as “Other”. We further grouped all non-coding RNAs into an “ncRNA” category as needed. Any peaks that contained none of the above annotations were categorized as “deep intergenic”.

#### CLIP Peak Enrichment Analysis

Peak enrichment analysis was performed on the annotated peak matrix by employing stringent filters based on the biological complexity (i.e. replicate counts) and normalized read counts. For each group, peaks were first filtered to exclude any peaks that were not present in at least half of the respective replicates. The BC filtered peaks for each group were further filtered by the read counts threshold, which was set as the median of the summed normalized read counts.

From the filtered peaks, peak enrichment score was calculated by directly comparing the summed normalized read counts per peak in different groups. To avoid division by zero in enrichment score calculation, pseudocount (non-zero minimum normalized read counts across all samples) was added to all sample normalized read counts.

#### Metagene Analysis

Transcription start site, start codon, stop codon, transcription termination site, 5’ splice site, and 3’ splice site coordinates were extracted from Ensembl release 110 gtf file from GRCh38.

Extracted feature site coordinates were extended by 500 nucleotides upstream and downstream, and divided into 20 nucleotides long bins. Peak density in each bin was calculated by counting the appearance of peaks in the bin and normalizing the counts by the total peak counts.

#### Sequence Motif Analysis

Respective filtered peaks for fractionation, input, and CoCLIP samples were used for sequence motif analysis. Using the peak coordinates, 150nt long sequences centered at the peak centers were extracted using the R BSgenome package and appearance of each HuR motif was counted for motif enrichment analysis. For HuR eCLIP, a peak file (ENCFF566LNK) was downloaded from the ENCODE consortium.^63^ Motif counts were then used to rank order the motifs for comparison across different samples. For de novo motif enrichment and motif density analysis, all mapped reads for respective groups were combined and used as input for HOMER to call separate peaks as described above, Then, findMotifsGenome and annotatePeaks were used with settings recommended in the program documentation to calculate motif enrichment and motif density near peaks. Finally, peaks were classified by the number of AAAAA or UUUUU motifs they contained and the distribution of the number of peaks per motif occurrences were normalized to generate cumulative distributions.

#### RNASeq Analysis

Sequencing FASTQ files were quality filtered and adapter trimmed using trim-galore software (https://github.com/FelixKrueger/TrimGalore) with default settings for paired-end sequencing data. Processed reads were then mapped to the human Hg38 reference genome using the *Rsubread subjunc*^64^ program allowing for a maximum of 3 mismatches in the reads. Gene level counts information was generated using *featureCounts* with the GENCODE v43 gtf file from GRCh38. Differential gene expression analysis was performed by following the standard *DESeq2* analysis protocol.^65^

#### Transcript Lengths Analysis

For genes that qualified for our location and condition specific analysis, transcript lengths and support level information were downloaded using Ensembl BioMart. For each gene, transcripts with support level of 1 or NA were kept as the representative gene lengths.

### Quantification and statistical analysis

All statistical analyses were performed in R using the stats package. For cumulative distributions comparison in motif analysis, the Kolmogorov-Smirnov test (ks.test) was used. For comparisons of lengths of genes enriched in different compartments, Mann-Whitney test (wilcox.test) was used.

## Data and code availability

All the sequencing data generated by this study have been deposited in the NCBI Gene Expression Omnibus (GEO) database under accession numbers GSE245209 and GSE245210. All original code for analysis and data visualization is freely available on GitHub at https://github.com/LunaRNALab/CLIPittyClip and https://github.com/LunaRNALab/BITs.

## Notes

https://www.ncbi.nlm.nih.gov/geo/query/acc.cgi?acc=GSE245209

https://www.ncbi.nlm.nih.gov/geo/query/acc.cgi?acc=GSE245210

https://github.com/LunaRNALab/CLIPittyClip

https://github.com/LunaRNALab/BITs

### Competing Interest Statement

The authors have declared no competing interest.

