## Supplementary Data for "Mapping RNA-Protein Interactions with Subcellular Resolution Using Colocalization CLIP"

**Supplemental Items:**

Figure S1: Proximity labeling for CoCLIP in three subcellular regions (related to Figure 1)
Figure S2: CoCLIP optimization and autoradiograms (related to Figure 1)
Figure S3: Summary of a coCLIP data analysis pipeline with CLIPittyClip and BITs (related to Figure 1)
Figure S4: Nuclear and cytoplasmic fractionation for HuR-CLIP (related to Figure 2)
Figure S5: RNA-Seq correlation between samples (related to Figure 5)

**Supplementary Tables:**

Table S1: Mapping Statistics (related to Figure 1-4, S3)
Table S2: HOMER Output Summary (related to Figure 4)
Table S3: Oligonucleotides (related to Figures 1, 3, S1, S2)

**Figure S1**

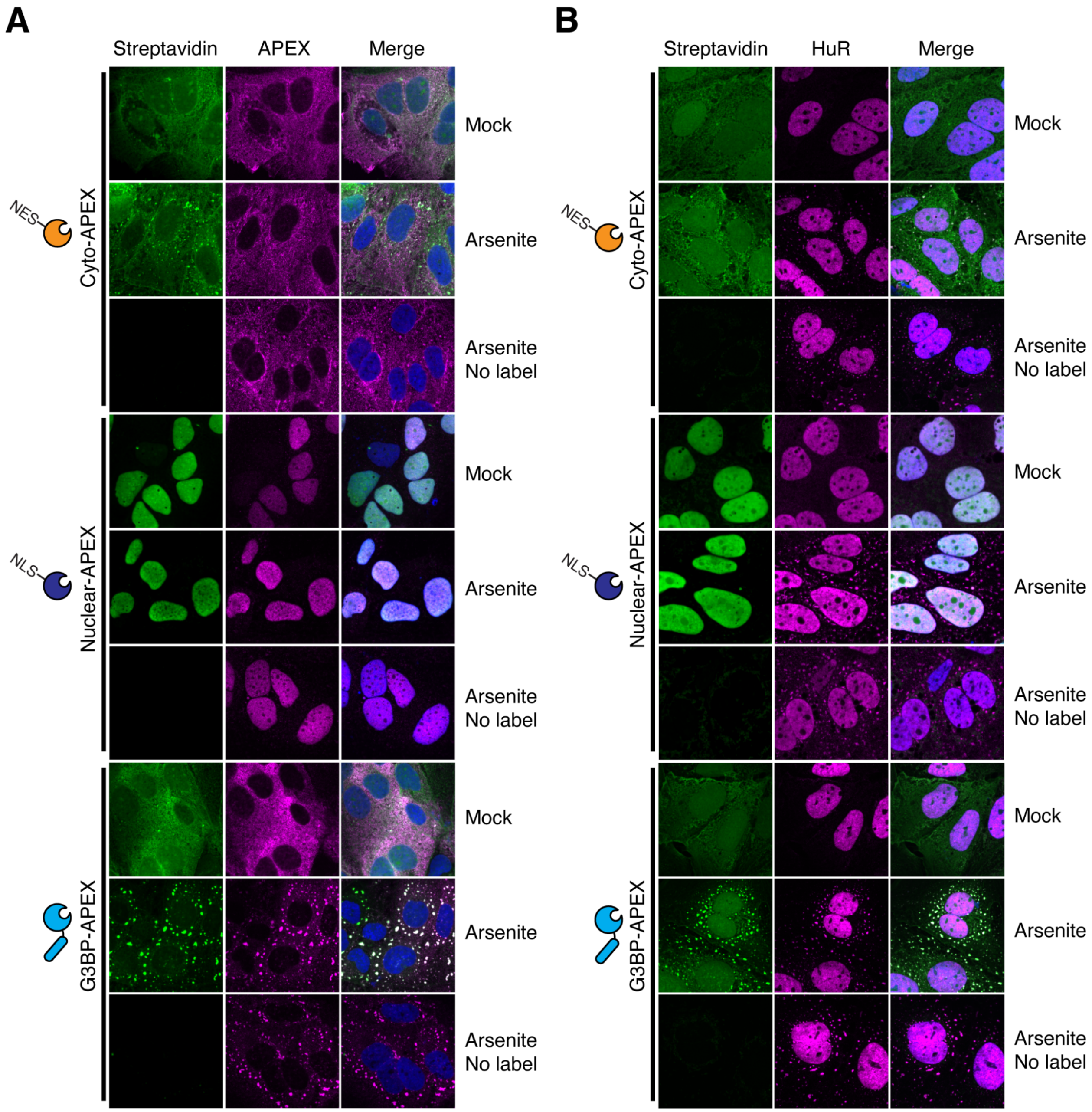


**Figure S1: Proximity labeling for CoCLIP in three subcellular regions. (A)** Immunofluorescence staining of Huh7.5 cells stably expressing APEX2 localized to the cytoplasm, nucleus or fused to the stress granule marker G3BP, in the presence or absence of arsenite stress. Fluorescent streptavidin was used to confirm site-specific labeling. **(B)** Immunofluorescence staining as in **(A)** showing HuR relocalization to stress granules upon arsenite stress for all conditions.

**Figure S2**

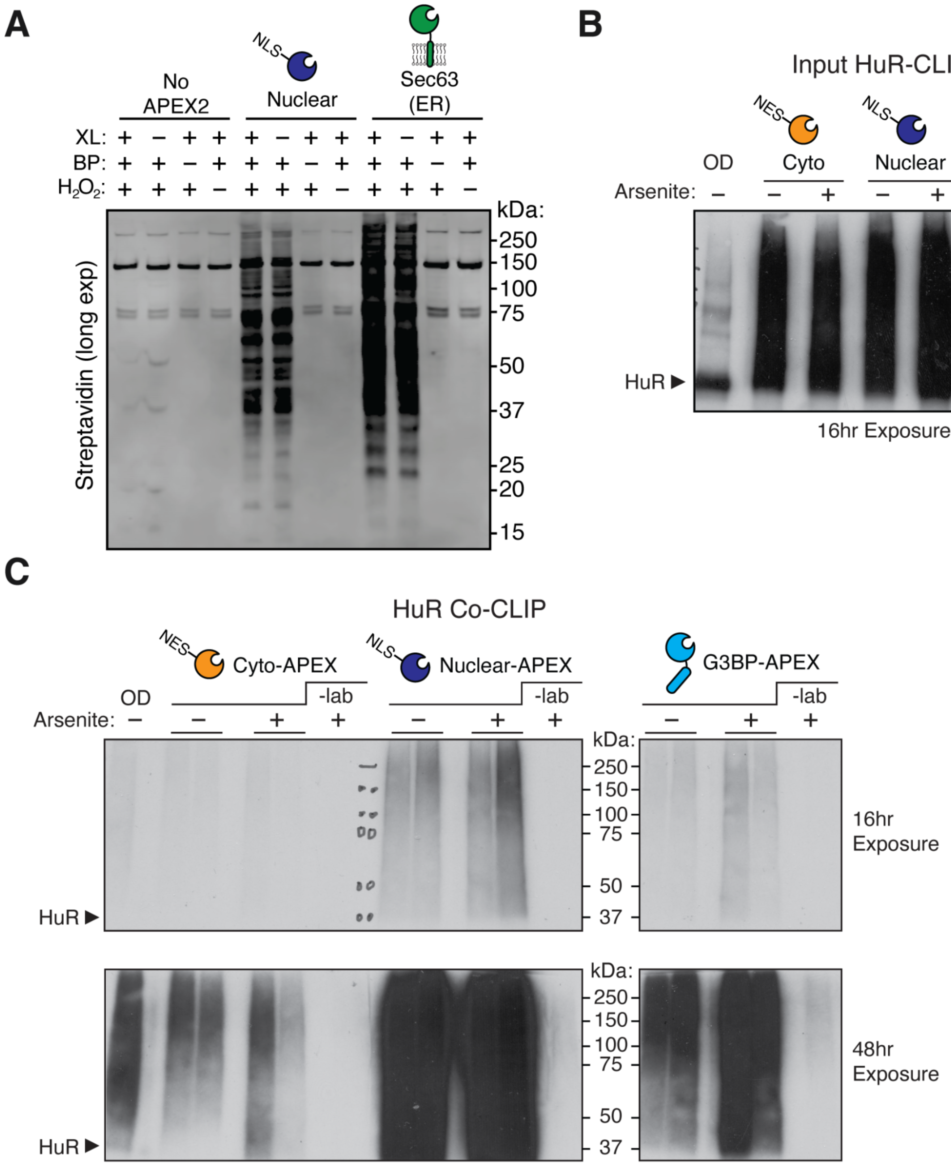


**Figure S2: CoCLIP optimization and autoradiograms. (A)** UV crosslinking does not induce spurious APEX2 activity. Immunoblot with fluorescent streptavidin in Huh7.5 cells with indicated APEX2 fusions, in the presence or absence of labeling reagents or UV crosslinking (XL) as indicated. BP: Biotin-Phenol. **(B)** Autoradiogram of total HuR CLIP from the conditions in Figure S1. OD: RNAse overdigest. **(C)** Autoradiogram of HuR coCLIP from the duplicate conditions in **(B)**, presented at two different exposures. Minus label control was performed by omitting hydrogen peroxide pulse.

#### Figure S3

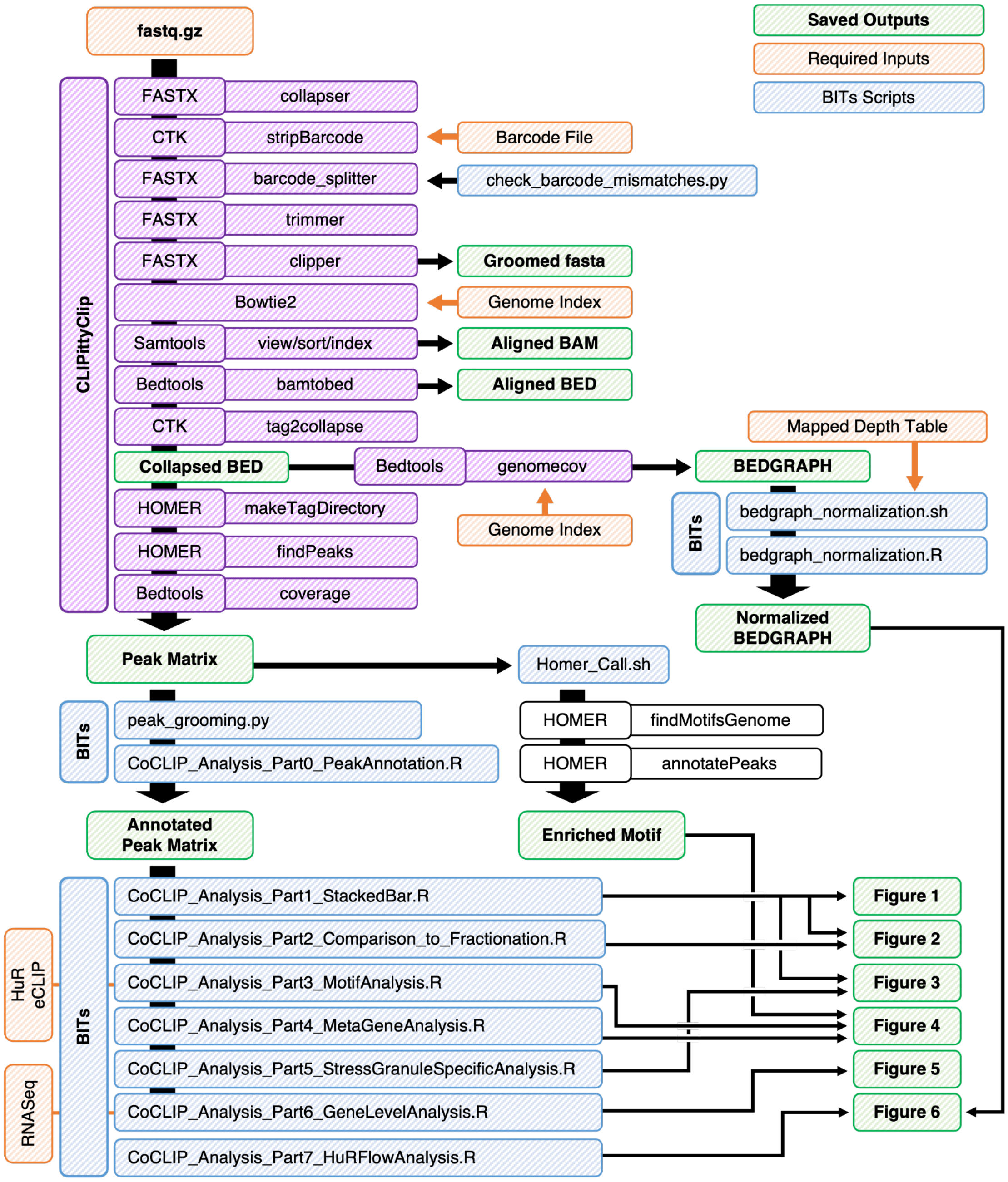


**Figure S3: Summary of a coCLIP data analysis pipeline with CLIPittyClip and BITs.** Flowchart diagram of data inputs, outputs, and ordered usage of tools used to process, analyze and visualize the data in this manuscript. CLIPittyClip provides a framework to process raw FASTQ files into a finished peak matrix in a single command line. BITs, for BioInformatics Tools and Scripts, enables downstream processing of CLIPittyClip outputs for visualization.

**Figure S4**

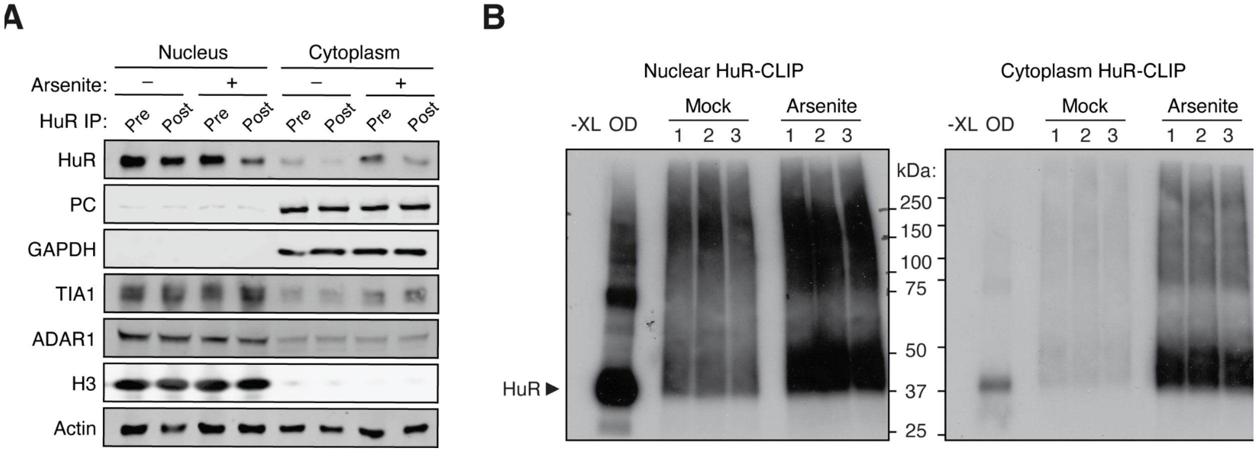


**Figure S4: Nuclear and cytoplasmic fractionation for HuR-CLIP. (A)** Western blot of nuclear or cytoplasmic fractions before and after HuR immunoprecipitation. Arsenite treatments as indicated. Pyruvate carboxylase (PC) and GAPDH were used as cytoplasmic markers; TIA1 and ADAR1 were used to mark the nucleoplasm; histone H3 was used to mark chromatin; Actin served as a general loading control. **(B)** Autoradiograms of HuR CLIP from nuclear (left) or cytoplasmic (right) fractions as in **(A)**. Exposure time was 30 minutes. -XL: no UV crosslinking control; OD: RNAse overdigest.

**Figure S5**

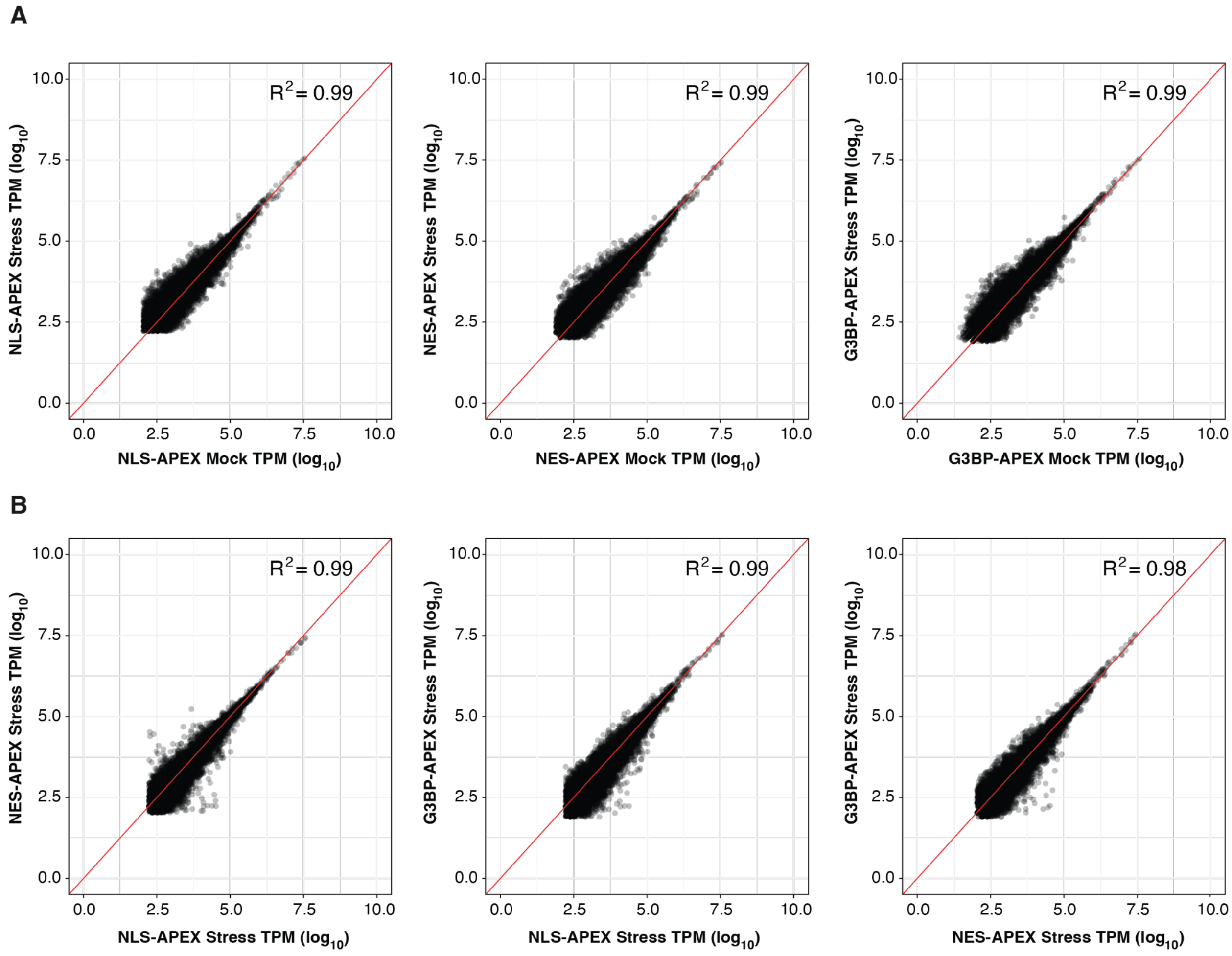


**Figure S5: RNA-Seq correlation between samples.** Pair-wise scatterplots of RNAseq transcript per million (TPM) measurements from APEX expressing cells used for coCLIP under mock **(A)** or arsenite **(B)** conditions. Pearson correlation coefficients are indicated.

#### Table S1: Mapping Statistics

| Data Source | Sample ID | Mapped Reads | Total Reads | % Mapped |
|---|---|---|---|---|
| JL0380_Nuc_Fraction_Mock_1 | Nuc_F_M_1 | 686263 | 703192 | 97.6 |
| JL0380_Nuc_Fraction_Mock_2 | Nuc_F_M_2 | 1099896 | 1124929 | 97.8 |
| JL0380_Nuc_Fraction_Mock_3 | Nuc_F_M_3 | 1035633 | 1059931 | 97.7 |
| JL0380_Nuc_Fraction_Ars_1 | Nuc_F_S_1 | 1248888 | 1277538 | 97.8 |
| JL0380_Nuc_Fraction_Ars_2 | Nuc_F_S_2 | 1250151 | 1280660 | 97.6 |
| JL0380_Nuc_Fraction_Ars_3 | Nuc_F_S_3 | 1154335 | 1180724 | 97.8 |
| JL0380_Cyto_Fraction_Mock_1 | Cyto_F_M_1 | 940147 | 963881 | 97.5 |
| JL0380_Cyto_Fraction_Mock_2 | Cyto_F_M_2 | 981792 | 1008995 | 97.3 |
| JL0380_Cyto_Fraction_Mock_3 | Cyto_F_M_3 | 948354 | 975909 | 97.2 |
| JL0380_Cyto_Fraction_Ars_1 | Cyto_F_S_1 | 1385165 | 1419657 | 97.6 |
| JL0380_Cyto_Fraction_Ars_2 | Cyto_F_S_2 | 1759090 | 1797001 | 97.9 |
| JL0380_Cyto_Fraction_Ars_3 | Cyto_F_S_3 | 1437680 | 1473264 | 97.6 |
| JL0388_NLS_Input_Mock_1 | NLS_I_M_1 | 1619414 | 1642052 | 98.6 |
| JL1024_NLS_Input_Mock_All | NLS_I_M_2 | 2622677 | 3061127 | 85.7 |
| JL0388_NLS_Input_Ars_1 | NLS_I_S_1 | 2295408 | 2317050 | 99.1 |
| JL1024_NLS_Input_Ars_All | NLS_I_S_2 | 1784078 | 2092738 | 85.3 |
| JL0388_NES_Input_Mock_1 | NES_I_M_1 | 834226 | 844647 | 98.8 |
| JL1024_NES_Input_Mock_All | NES_I_M_2 | 120962 | 163387 | 74 |
| JL0388_NES_Input_Ars_1 | NES_I_S_1 | 404260 | 413705 | 97.7 |
| JL1024_NES_Input_Ars_All | NES_I_S_2 | 122616 | 167305 | 73.3 |
| JL0361_G3BP_Input_Mock_1 | G3BP_I_M_1 | 19748 | 22417 | 88.1 |
| JL0361_G3BP_Input_Mock_2 | G3BP_I_M_2 | 210794 | 218943 | 96.3 |
| JL0388_G3BP_Input_Mock_1 | G3BP_I_M_3 | 765265 | 774742 | 98.8 |
| JL1024_G3BP_Input_Mock_All | G3BP_I_M_4 | 22476 | 89907 | 25 |
| JL0361_G3BP_Input_Ars_1 | G3BP_I_S_1 | 17051 | 19530 | 87.3 |
| JL0361_G3BP_Input_Ars_2 | G3BP_I_S_2 | 1109018 | 1146723 | 96.7 |
| JL0361_G3BP_Input_Ars_3 | G3BP_I_S_3 | 1206979 | 1239967 | 97.3 |
| JL0388_G3BP_Input_Ars_1 | G3BP_I_S_4 | 829192 | 838876 | 98.8 |
| JL1024_G3BP_Input_Ars_All | G3BP_I_S_5 | 180888 | 270815 | 66.8 |
| JL0388_NLS_Enrich_Mock_1 | NLS_E_M_1 | 202951 | 211314 | 96 |
| JL0388_NLS_Enrich_Mock_2 | NLS_E_M_2 | 146321 | 151918 | 96.3 |
| JL1024_NLS_Enrich_Mock_R1 | NLS_E_M_3 | 528719 | 661200 | 80 |
| JL1024_NLS_Enrich_Mock_R2 | NLS_E_M_4 | 477565 | 554969 | 86.1 |
| JL0388_NLS_Enrich_Ars_1 | NLS_E_S_1 | 72101 | 75122 | 96 |
| JL0388_NLS_Enrich_Ars_2 | NLS_E_S_2 | 224335 | 229473 | 97.8 |
| JL1024_NLS_Enrich_Ars_R1 | NLS_E_S_3 | 523287 | 651909 | 80.3 |
| JL1024_NLS_Enrich_Ars_R2 | NLS_E_S_4 | 344414 | 426354 | 80.8 |

| | | | | |
|---|---|---|---|---|
| JL0388_NES_Enrich_Mock_1 | NES_E_M_1 | 37831 | 44483 | 85 |
| JL0388_NES_Enrich_Mock_2 | NES_E_M_2 | 50343 | 54651 | 92.1 |
| JL1024_NES_Enrich_Mock_R1 | NES_E_M_3 | 114360 | 196682 | 58.1 |
| JL1024_NES_Enrich_Mock_R2 | NES_E_M_4 | 105485 | 198586 | 53.1 |
| JL0388_NES_Enrich_Ars_1 | NES_E_S_1 | 38255 | 44650 | 85.7 |
| JL0388_NES_Enrich_Ars_2 | NES_E_S_2 | 76213 | 80852 | 94.3 |
| JL1024_NES_Enrich_Ars_R1 | NES_E_S_3 | 482900 | 656758 | 73.5 |
| JL1024_NES_Enrich_Ars_R2 | NES_E_S_4 | 564779 | 777109 | 72.7 |
| JL0361_G3BP_Enrich_Mock_1 | G3BP_E_M_1 | 6330 | 11268 | 56.2 |
| JL0361_G3BP_Enrich_Mock_2 | G3BP_E_M_2 | 151280 | 175806 | 86 |
| JL0388_G3BP_Enrich_Mock_1 | G3BP_E_M_3 | 14760 | 19822 | 74.5 |
| JL0388_G3BP_Enrich_Mock_2 | G3BP_E_M_4 | 4662 | 9028 | 51.6 |
| JL1024_G3BP_Enrich_Mock_R1 | G3BP_E_M_5 | 334236 | 564930 | 59.2 |
| JL1024_G3BP_Enrich_Mock_R2 | G3BP_E_M_6 | 944225 | 1200994 | 78.6 |
| JL0361_G3BP_Enrich_Ars_1 | G3BP_E_S_1 | 13149 | 18848 | 69.8 |
| JL0361_G3BP_Enrich_Ars_2 | G3BP_E_S_2 | 604651 | 741496 | 81.5 |
| JL0361_G3BP_Enrich_Ars_3 | G3BP_E_S_3 | 524332 | 642651 | 81.6 |
| JL0388_G3BP_Enrich_Ars_1 | G3BP_E_S_4 | 88707 | 91738 | 96.7 |
| JL0388_G3BP_Enrich_Ars_2 | G3BP_E_S_5 | 28062 | 31480 | 89.1 |
| JL1024_G3BP_Enrich_Ars_R1 | G3BP_E_S_6 | 117098 | 148123 | 79.1 |
| JL1024_G3BP_Enrich_Ars_R2 | G3BP_E_S_7 | 66951 | 106854 | 62.7 |

### Table S2: HOMER Output Summary

* possible false-positive reported by HOMER

**All Libraries:**

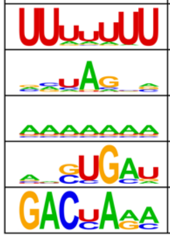

| Motif | p-value |
|---|---|
| | 1e-11810 |
| | 1e-1651 |
| | 1e-275 |
| | 1e-254 |
| | 1e-171 |

**CoCLIP Input Mock:**

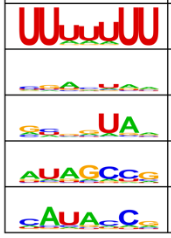

| Motif | p-value |
|---|---|
| | 1e-1568 |
| | 1e-145 |
| | 1e-107 |
| | 1e-42 |
| | 1e-37 |

**CoCLIP Input Arsenite:**

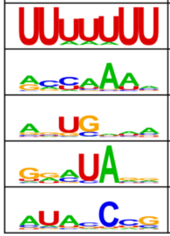

| Motif | p-value |
|---|---|
| | 1e-2427 |
| | 1e-274 |
| | 1e-241 |
| | 1e-121 |
| | 1e-55 |

**CoCLIP NLS Mock:**

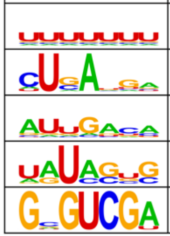

| Motif | p-value |
|---|---|
| | 1e-95 |
| | 1e-10* |
| | 1e-10* |
| | 1e-10* |
| | 1e-9* |

**CoCLIP NLS Arsenite:**

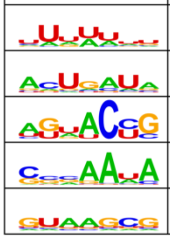

| Motif | p-value |
|---|---|
| | 1e-132 |
| | 1e-13 |
| | 1e-11* |
| | 1e-10* |
| | 1e-9* |

**CoCLIP NES Mock:**

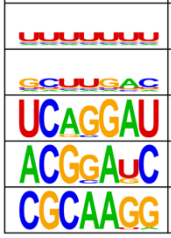

| Motif | p-value |
|---|---|
| | 1e-17 |
| | 1e-17 |
| | 1e-11* |
| | 1e-9* |
| | 1e-8* |

**CoCLIP NES Arsenite:**

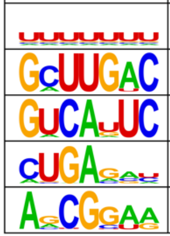

| Motif | p-value |
|---|---|
| | 1e-64 |
| | 1e-13 |
| | 1e-10* |
| | 1e-10* |
| | 1e-8* |

**CoCLIP G3BP Mock:**

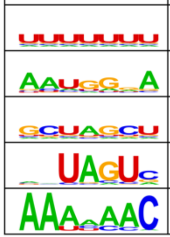

| Motif | p-value |
|---|---|
| | 1e-573 |
| | 1e-10* |
| | 1e-9* |
| | 1e-8* |
| | 1e-8* |

**CoCLIP G3BP Arsenite:**

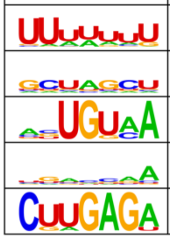

| Motif | p-value |
|---|---|
| | 1e-274 |
| | 1e-14 |
| | 1e-13 |
| | 1e-13 |
| | 1e-13 |

**Nuclear Fraction CLIP Mock:**

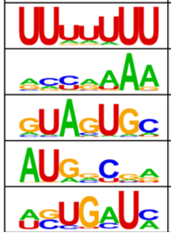

| Motif | p-value |
|---|---|
| | 1e-3553 |
| | 1e-362 |
| | 1e-89 |
| | 1e-86 |
| | 1e-54 |

**Nuclear Fraction CLIP Arsenite:**

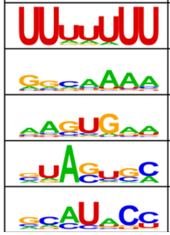

| Motif | p-value |
|---|---|
| | 1e-3964 |
| | 1e-262 |
| | 1e-121 |
| | 1e-56 |
| | 1e-48 |

**Cytoplasm Fraction CLIP Mock:**

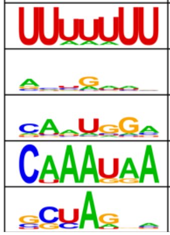

| Motif | p-value |
|---|---|
| | 1e-2838 |
| | 1e-348 |
| | 1e-56 |
| | 1e-44 |
| | 1e-43 |

**Cytoplasm Fraction CLIP Arsenite:**

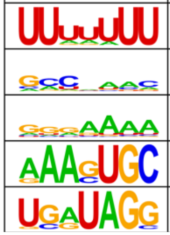

| Motif | p- |
|---|---|
| | 1e-5355 |
| | 1e-515 |
| | 1e-362 |
| | 1e-62 |
| | 1e-36 |

#### Table S3: Oligonucleotides

| PCR Primers | | | |
|---|---|---|---|
| **Name** | **Sequence** | **Content** | **Primer partner** |
| oJL529 | GCTCGGTACCCGGGGATC**ATCGAT**GCCACCATGGACTACAAGGATGACG | ePB_ClaI_APEX2_F | 530 |
| oJL578 | GTACCCGGGGATC**ATCGAT**GCCACCATGGGCCCTAAAAAGAAGCGTAAAGTCGACTACAAGGATGACGACGATAAGG | ePB_ClaI_nls-APEX2_F | 530 |
| oJL579 | GTACCCGGGGATC**ATCGAT**GCCACCATGTTAGCTTTGAAATTAGCCGGACTAGACATCGACTACAAGGATGACGACGATAAGG | ePB_ClaI_nes-APEX2_F | 530 |
| oJL530 | CTTAATCAGCTCGCTCACCATTCC**GGATCC**GGCATCAGCAAACCCAAGCT | mKate2_BamHI_APEX2_R | 529, 578, 579 |
| oJL531 | AGCTTGGGTTTGCTGATGCC**GGATCC**GGAATGGTGAGCGAGCTGATTAAG | APEX2_BamHI_mKate2_F | 532 |
| oJL532 | GGCTGATTATGATCTAGAGTC**GCGGCCGC**TCTGTGCCCCAGTTTGCTAG | ePB_NotI_mKate2_R | 531 |
| oJL618 | GCAAACTGGGGCACAGA**GCTAGC**GGAGTGATGGAGAAGCCTAGTCCC | mKate2_NheI_G3BP1_F | 619 |
| oJL619 | CAGGTTCCTCTACTTCTTCTTCCTCCTCCTGAGGCTCAGTGACAA | G3BP1_S149E_R | 618 |
| oJL620 | TTGTCACTGAGCCTCAGGAGGAGGAAGAAGAAGAAGTAGAGGAACCTG | G3BP1_S149E_F | 516 |
| oJL516 | GGCTGATTATGATCTAGAGTC**GCGGCCGC**TTACTGCCGTGGCGCAAG | ePB_NotI_G3BP1_R | 620 |
| oJL573 | GTACCCGGGGATC**ATCGAT**GCCACCATGGCCGGGCAGCAGTTCC | ePB-Sec63_F | 577 |
| oJL577 | TCCTTAATCAGCTCGCTCACCAT**CGGCCG**ACGATACCACATACACCTTCCATATA | mKate2_EagI_Sec63_R | 573 |
| oJL578 | TATATGGAAGGTGTATGTGGTATCGT**CGGCCG**ATGGTGAGCGAGCTGATTAAGGA | Sec63_EagI_mKate2_F | 574 |

| | | | |
|---|---|---|---|
| oJL574 | CGTCGTCATCCTTGTAGTCCATTCC**GGATCC**TCTGTGCCCCAGTTTGCTAGGG | APEX2_BamHI_mKate2_R | 578 |
| oJL575 | CTAGCAAACTGGGGCACAGA**GGATCC**GGAATGGACTACAAGGATGACGACGATAA | mKate2_BamHI_APEX2_F | 576 |
| oJL576 | GGCTGATTATGATCTAGAGTC**GCGGCCGC**TTAGGCATCAGCAAACCCAAGCT | ePB_NotI_APEX2_R | 575 |
| **CLIP Oligos** | | | |
| **Name** | **Sequence** | **Type** | **Index** |
| L32 | /5Phos/GUGUCAGUCACUUCCAGCGG/3ddC/ | CLIP RNA linker | N/A |
| RT-1T | /5Phos/DDDCGTGATNNNNNNNAGATCGGAAGAGCGTCGT/iSp18/CACTCA/iSp18/CCGCTGGAAGTGACTGAC | CLIP RT Oligos | ATCACG |
| RT-2T | /5Phos/DDDACATCGNNNNNNNAGATCGGAAGAGCGTCGT/iSp18/CACTCA/iSp18/CCGCTGGAAGTGACTGAC | | CGATGT |
| RT-3T | /5Phos/DDDGCCTAANNNNNNNAGATCGGAAGAGCGTCGT/iSp18/CACTCA/iSp18/CCGCTGGAAGTGACTGAC | | TTAGGC |
| RT-4T | /5Phos/DDDTGGTCANNNNNNNAGATCGGAAGAGCGTCGT/iSp18/CACTCA/iSp18/CCGCTGGAAGTGACTGAC | | TGACCA |
| RT-5T | /5Phos/DDDCACTGTNNNNNNNAGATCGGAAGAGCGTCGT/iSp18/CACTCA/iSp18/CCGCTGGAAGTGACTGAC | | ACAGTG |
| RT-6T | /5Phos/DDDATTGGCNNNNNNNAGATCGGAAGAGCGTCGT/iSp18/CACTCA/iSp18/CCGCTGGAAGTGACTGAC | | GCCAAT |
| RT-7T | /5Phos/DDDGATCTGNNNNNNNAGATCGGAAGAGCGTCGT/iSp18/CACTCA/iSp18/CCGCTGGAAGTGACTGAC | | CAGATC |
| RT-8T | /5Phos/DDDTCAAGTNNNNNNNAGATCGGAAGAGCGTCGT/iSp18/CACTCA/iSp18/CCGCTGGAAGTGACTGAC | | ACTTGA |
| RT-9T | /5Phos/DDDCTGATCNNNNNNNAGATCGGAAGAGCGTCGT/iSp18/CACTCA/iSp18/CCGCTGGAAGTGACTGAC | | GATCAG |
| RT-10T | /5Phos/DDDAAGCTANNNNNNNAGATCGGAAGAGCGTCGT/iSp18/CACTCA/iSp18/CCGCTGGAAGTGACTGAC | | TAGCTT |
| RT-11T | /5Phos/DDDGTAGCCNNNNNNNAGATCGGAAGAGCGTCGT/iSp18/CACTCA/iSp18/CCGCTGGAAGTGACTGAC | | GGCTAC |
| RT-12T | /5Phos/DDDTACAAGNNNNNNNAGATCGGAAGAGCGTCGT/iSp18/CACTCA/iSp18/CCGCTGGAAGTGACTGAC | | CTTGTA |

| | | | |
|---|---|---|---|
| RT-13T | /5Phos/DDDCGTGATNNNNNNNAGATCGGAAGAGCGTCGT/iSp18/CACTCA/iSp18/CCGCTGGAAGTGACTGAC | | ATCACG |
| RT-14T | /5Phos/DDDGGAACTNNNNNNNAGATCGGAAGAGCGTCGT/iSp18/CACTCA/iSp18/CCGCTGGAAGTGACTGAC | | AGTTCC |
| RT-15T | /5Phos/DDDTGACATNNNNNNNAGATCGGAAGAGCGTCGT/iSp18/CACTCA/iSp18/CCGCTGGAAGTGACTGAC | | ATGTCA |
| RT-16T | /5Phos/DDDGGACGGNNNNNNNAGATCGGAAGAGCGTCGT/iSp18/CACTCA/iSp18/CCGCTGGAAGTGACTGAC | | CCGTCC |
| RT-18T | /5Phos/DDDGCGGACNNNNNNNAGATCGGAAGAGCGTCGT/iSp18/CACTCA/iSp18/CCGCTGGAAGTGACTGAC | | GTCCGC |
| RT-19T | /5Phos/DDDTTTCACNNNNNNNAGATCGGAAGAGCGTCGT/iSp18/CACTCA/iSp18/CCGCTGGAAGTGACTGAC | | GTGAAA |
| RT-20T | /5Phos/DDDGGCCACNNNNNNNAGATCGGAAGAGCGTCGT/iSp18/CACTCA/iSp18/CCGCTGGAAGTGACTGAC | | GTGGCC |
| RT-21T | /5Phos/DDDCGAAACNNNNNNNAGATCGGAAGAGCGTCGT/iSp18/CACTCA/iSp18/CCGCTGGAAGTGACTGAC | | GTTTCG |
| DP5 | AATGATACGGCGACCACCGAGATCTACACTCTTTCCCTACACGACGCTCTTCCGATCT | CLIP PCR Primers | N/A |
| SP3 | CAAGCAGAAGACGGCATACGAGATCTCGGCATTCCTGCCGCTGGAAGTGACTGACAC | | N/A |
